# Nanoparticle size governs engagement with myeloid cells and hitchhiking to hematopoietic organs in myeloproliferative neoplasms

**DOI:** 10.64898/2026.09.07.749819

**Authors:** Sara Elsafy, Shiva Khorshid, Alessia Nucci, Federica De Lorenzi, Alessandro Motta, Margherita Vieri, Jelena Lazarevic, Mithuoshni Arullmoli, Julian Baumeister, Nicolas Chatain, Eva Miriam Buhl, Fabian Kiessling, Twan Lammers, Steffen Koschmieder, Josbert M. Metselaar, Marcelo Augusto Szymanski de Toledo, Alexandros Marios Sofias

**Affiliations:** Institute for Experimental Molecular Imaging (ExMI), RWTH Aachen University Hospital, Aachen, Germany; Department of Hematology, Oncology, Hemostaseology and Stem Cell Transplantation, Faculty of Medicine, RWTH Aachen University Hospital, Aachen, Germany; Center for Integrated Oncology Aachen (CIOA), RWTH Aachen University Hospital, Aachen, Germany; Electron Microscopy Facility, Institute of Pathology, University Hospital Aachen, RWTH Aachen University, Aachen, Germany

**Keywords:** Nanomedicine, continuous flow manufacturing, liposomes, myeloid immune cells, neutrophils, bone marrow targeting, myeloproliferative neoplasms (MPN), image-guided drug delivery

## Abstract

Nanoparticle design principles contribute to desirable *in vivo* performance, including prolonged circulation, desirable biodistribution profiles, and tuned interactions with the immune system. The importance of these variables, although well-established in solid tumors, remains elusive in the context of hematological malignancies. Here, we investigated the influence of liposome size on biodistribution and cellular uptake within hematopoietic compartments – bone marrow (BM) and spleen – in *JAK2*^V617F^ myeloproliferative neoplasms (MPN). A milli-fluidic manufacturing platform combined with a Design-of-Experiments (DoE) approach was used to generate small, medium, and large liposomes. Liposome uptake was assessed *ex vivo* using blood samples from healthy donors and MPN patients, followed by *in vivo* biodistribution studies in a transgenic *JAK2*^V617F^ MPN mouse model. Organ-level accumulation was quantified using hybrid fluorescence / computed tomography (FLT/CT) imaging, while cellular uptake was evaluated via flow cytometry. Whole-body imaging revealed that increasing liposome size enhanced delivery to both the spleen and BM, with larger liposomes exhibiting the highest accumulation in both organs. Cellular analysis corroborated the *in vivo* observations, demonstrating that large liposomes are taken up to a greater extent by monocytes and granulocytes, in both spleen and BM. At late time-points post-injection, the liposomal accumulation was twice as high in the BM in comparison to the spleen and peripheral blood, highlighting the progressively elevated accumulation and retention in the BM, as opposed to the elimination phase nanoparticles undergo in clearance organs and circulation at these time-points. Of note, among BM mature myeloid cells, neutrophils were associated with a higher nanoparticle uptake than monocytes, highlighting their capability to phagocytose material in circulation and hitchhike it to malignant or inflamed regions. These findings demonstrate that continuous flow manufacturing procedures can swiftly produce nanoparticles with desirable characteristics that are essential for tuning the biodistribution towards hematopoietic organs and myeloid immune cells. By providing mechanistic insights into nanoparticle behavior within hematopoietic compartments, this work advances our understanding of nanomedicine *in vivo* performance and highlights particle size as a critical quality attribute that can be optimized and controlled to improve targeted delivery to myeloid cells in the treatment of hematological malignancies.

**Graphical abstract:** 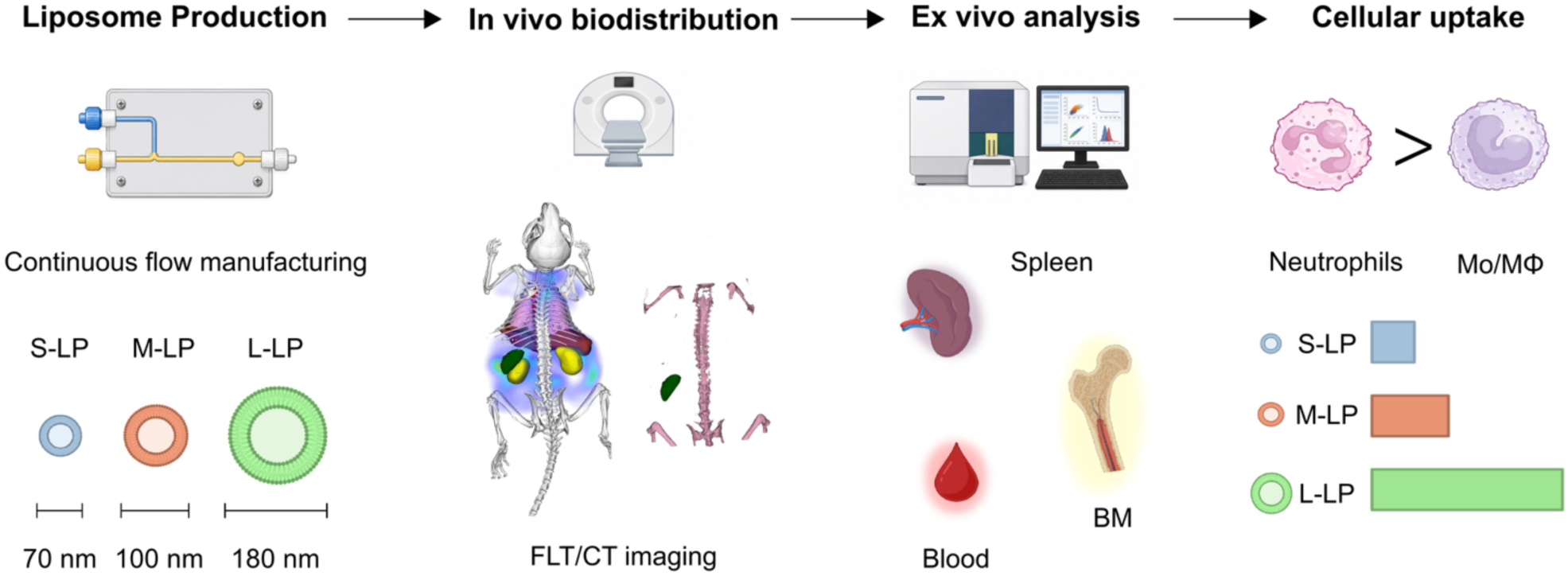

## 1. Introduction

Nanomedicines improve drug delivery by enhancing accumulation at disease-relevant sites while reducing systemic exposure^1–5^. Most nanoparticle design principles stem from solid tumor delivery studies and rely on the notion that abnormal tumor vasculature containing inter-endothelial gaps permits small nanocarriers to extravasate into tumor tissue and be retained due to impaired lymphatic drainage^4,6–10^, thereby favoring particles around 100 nm, as they provide an optimal balance between prolonged circulation and efficient tumor accumulation^3,4,9,11,12^.

Unlike solid tumors, hematological malignancies arise within hematopoietic tissues, especially the bone marrow (BM), disseminate through the circulation, and establish malignant lesions at the systemic level. The hematopoietic tissues possess distinct vascular and cellular characteristics that likely impose different constraints on nanoparticle transport, accumulation, and cellular uptake^1,13–17^. In addition, unlike solid malignancies, hematopoietic tissues contain larger numbers of myeloid cells and undergo rapid cellular turnover. While interactions of nanoparticles with myeloid immune cells have traditionally been regarded as a barrier to effective drug delivery, growing evidence suggests that they provide opportunities for targeted delivery and therapy^18–24^. These observations suggest that the size requirements governing nanoparticle accumulation and cellular uptake in hematopoietic tissues may differ substantially from those established in solid tumors. Nevertheless, the physicochemical determinants of nanoparticle distribution within hematopoietic tissues remain poorly understood.

To better understand nanoparticle size behavior within hematopoietic compartments, myeloproliferative neoplasms (MPN) were selected as a disease model. MPN are a spectrum of chronic hematological malignancies characterized by the clonal expansion of myeloid lineage cells in the BM. It comprises three major phenotypes, namely essential thrombocythemia (ET), polycythemia vera (PV), and primary myelofibrosis (PMF). These MPN subtypes share common primary driver mutations in Janus kinase 2 (*JAK2*), Calreticulin (*CALR*), or Myeloproliferative Leukemia Virus Oncogene (*MPL*) genes, leading to constitutive JAK2/STAT activation and uncontrolled myeloid cell proliferation^25–27^. In addition to BM pathology, MPN are frequently associated with splenomegaly and expansion of myeloid populations across the spleen and peripheral blood. These hematopoietic compartments remain interconnected through continuous myeloid cell trafficking. Notably, myeloid cell trafficking is not restricted to unidirectional egress from the BM into the circulation but can also involve recirculation back to the BM in response to homeostatic and inflammatory cues^28–34^. Such bidirectional trafficking may potentially influence nanoparticle accumulation between hematopoietic compartments. Together, these features make MPN a suitable model for studying nanoparticle interactions within the hematopoietic organs and for potentially identifying new opportunities for therapeutic delivery to these sites.

Liposomes were chosen among other delivery systems, as they are a major class of clinically approved nanomedicines and have been widely used to deliver anticancer therapeutics^5,25,35,36^. Considering their preparation methods, conventional batch-to-batch protocols, although well-established, often require post-processing manufacturing steps that are difficult to control in terms of batch reproducibility and during scale-up, and frequently result in heterogeneous size distributions^35^. To this end, continuous flow manufacturing approaches, including milli-fluidic systems, with in-flow process analytical technology (PAT) allow precise control over nanoparticle size and physicochemical properties. By regulating parameters such as lipid concentration, total flow rate (TFR), and flow rate ratio (FRR), these systems enable controlled mixing of aqueous and organic phases and thereby achieve more reproducible production of liposomes with defined sizes^37–43^. Importantly, such approaches support translational development because continuous flow manufacturing controlled by PAT offers scalability^44^.

Taken together, in this study, we optimized a milli-fluidic manufacturing platform using a Design-of-Experiments (DoE) approach to systematically investigate the effect of manufacturing parameters on liposome size. Subsequently, we identified and selected conditions that enable reproducible production of liposomes with defined sizes: small liposomes (S-LP), 70 nm; medium liposomes (M- LP), 100 nm; and large liposomes (L-LP), 180 nm. We then evaluated the influence of particle size on monocyte and granulocyte uptake using *ex vivo* whole blood samples obtained from healthy donors and MPN patients, followed by *in vivo* biodistribution studies in a transgenic *JAK2*^V617F^ MPN mouse model that recapitulates key features of PV. Liposome accumulation was assessed at both the organ level using hybrid fluorescence / computed tomography (FLT/CT) imaging and at the cellular level using flow cytometry, with a particular focus on hematopoietic compartments such as the BM and spleen. Through this approach, we aimed to identify size-dependent distribution patterns that may inform the rational design of nanomedicines for hematologic malignancies.

## 2. Materials and methods

### 2.1. Materials

DPPC 16:0 PC (1,2-dipalmitoyl-sn-glycero-3-phosphocholine) and mPEG-(2K)-DSPE 18:0 PE (N- (carbonyl-methoxypolyethylene glycol-2000) -1,2-distearoyl-sn-glycero-3-phosphoethanolamine) were obtained from Lipoid GmbH (Ludwigshafen, Germany). Cholesterol was acquired from Sigma-Aldrich (Merck, Germany). Cyanine 7 PE1,2-distearoyl-sn-glycero-3-phosphoethanolamine-N- (Cyanine 7) was purchased from Avanti^®^ Polar Lipids (Merck KGaA, Germany). DiD:DiIC18 (5) solid (1,1’-Dioctadecyl-3,3,3’,3’-Tetramethylindodicarbocyanine, 4-Chlorobenzenesulfonate Salt) dye was acquired from Invitrogen^™^ (Fisher Scientific, Germany). Spectra/Por^®^ regenerated cellulose dialysis membrane tubing (MWCO 12-14 kDa) was obtained from Spectrum^™^ Labs (Fisher Scientific, Germany). Phosphate buffered saline (PBS) ready-to-use tablets (ROTI^®^Fair PBS 7.4) and ethanol were purchased from Carl Roth GmbH (Karlsruhe, Germany). Amicon^®^ Ultra Centrifugal filter (10 kDa MWCO) and was obtained from Merck Millipore (Merck KGaA, Germany). Cholesterol FS Assay and cholesterol standard were purchased from DiaSys (Diagnostic Systems GmbH, Germany). Cell culture medium and supplements were obtained from Gibco (Thermo Fisher Scientific, Germany). EASYstrainer^™^ 40 µm and 70 µm cell strainers were bought from Greiner (Bio-One GmbH, Germany). Element HT5 Diluent, Element HT5 LH lyse, and Element HT5 Diff lysis were obtained from Medical Solution GmbH (Wil SG, Switzerland). Erythrocyte lysis buffer was purchased from Morphisto GmbH (Offenbach Germany). EDTA-blood sampling tubes were purchased from Sarstedt (Nümbrecht, Germany).

### 2.2. Design-of-Experiments

A DoE approach was employed to systematically evaluate the influence of formulation and process parameters on liposome (LP) physicochemical properties. Lipid concentration was assessed as a formulation parameter, while FRR and TFR were investigated as process parameters. Operating temperature was evaluated separately in a preliminary screening experiment and subsequently fixed at 70 °C for all DoE runs. Lipid concentrations ranged from 10 to 150 mM (total lipids dissolved in ethanol), FRRs were set at 1, 1.5, 2, and 2.5, and TFR varied from 40 to 140 ml/min. The effects of these parameters on LP size (Z-avg) and polydispersity index (PDI) were investigated. Design- Expert^®^ software (version 23.1, Stat-Ease Inc., USA) was used to generate the experimental matrix. A total of 45 runs across three blocks were performed using a D-optimal design based on a linear model complexity of the initial model, allowing the evaluation of multiple composition and process parameters and their interactions, while minimizing the number of experiments required. The detailed list of experimental conditions is provided in **Table S1**. In each run LP size and PDI were measured by dynamic light scattering (DLS) using a Zetasizer (Malvern Instruments Ltd., UK). A response surface model was composed to identify significant factors and interactions, and to generate predictive models. Non-significant terms (p > 0.05) were excluded from the model, except where needed to preserve model hierarchy. The software was then used to visualize the relationship between input variables and output responses through surface plots. The model’s predictive accuracy was evaluated by analyzing a scatterplot of predicted R2 (R2-pred) against experimental values and by calculating the correlation coefficient (R2).

### 2.3. Liposome preparation and characterization

LP were prepared using a continuous flow milli-fluidic device^44,45^. Briefly, a lipid solution was prepared by dissolving DPPC, cholesterol, and DSPE-PEG2000 in ethanol at a fixed molar ratio of 62:33:5, respectively. The aqueous phase used was PBS. Unless otherwise stated, both the lipid and aqueous solutions were preheated to 70 °C (above DPPC transition temperature) in a water bath. Once heated, they were transferred to syringes and loaded onto advanced programmable PHD ULTRA^™^ syringe pumps (Harvard Apparatus, USA). LP were formed by mixing the lipid and aqueous phases within the milli-fluidic device. For the DoE, lipid concentration, TFR, and FRR were varied as experimental factors. The specific values used are detailed in **Table S1**. The size and PDI of LP were measured using DLS on a Zetasizer Nano ZS90 (Malvern Instruments Ltd., UK). The LP were diluted 10x in PBS and measured at 25 °C after a 120-second equilibration period. Each sample was measured in triplicate, with the number of runs automatically determined by the instrument. Formed LP was dialyzed against PBS (pH 7.4) using Spectra/Por^®^ regenerated cellulose dialysis membranes (MWCO 12–14 kDa) with gentle stirring at 200 rpm for 24 hours to remove residual ethanol. LP were then stored at 4 °C until further use.

### 2.4. Preparation of liposomes for *in vivo* biodistribution

To generate LP of different sizes for *in vivo* experiments, FRR was varied between 1.5 and 2.5, while the temperature, initial lipid concentration, and TFR were fixed at 70°C, 20 mM, and 140 ml/min, respectively. The LP contained 0.2 mol% DSPE-Cy7 (Avanti® Polar Lipids, Merck KGaA, Germany) and had a lipid composition of DPPC:cholesterol:DSPE-PEG2000:DSPE-Cy7 with molar ratios of 62:33:4.8:0.2. Following dialysis, the LP were concentrated to a final lipid concentration of 10 mM using Amicon^®^ Ultra centrifugal filters. Briefly, 2 ml of LP suspension was transferred to the filter unit and centrifuged at 3,000 g for 40 minutes, or until the required lipid concentration was achieved. Concentrated LP were stored at 4 °C until further use.

### 2.5. Cryogenic transmission electron microscopy (Cryo-TEM)

Different-sized Cy7-labeled LPs were prepared as described in section 2.4. Cryo-TEM was used to assess the morphology of LP. Formulations were applied on Lacey Formvar/Carbon Nickel-grids (Plano, Germany) and plunge frozen in ethane using a plunge cooler (MiTeGen, USA). Vitrified samples were imaged using a HT7800 cryo-TEM (Hitachi, Japan) at 120 kV.

### 2.6. Lipid content quantification

To ensure equal lipid dosing across size-defined formulations, cholesterol content was quantified using the DiaSys Cholesterol FS assay (Diagnostic Systems GmbH, Germany), an enzymatic colorimetric assay. Briefly, a serial dilution of cholesterol standard was prepared in PBS, in triplicate, and placed in a transparent 96-well plate. Then 10 µl of each LP sample was added in triplicate to the 96-well plate. Next, 190 µl of the cholesterol reagent was added to both the standard curve wells and the LP sample wells. The 96-well plate was incubated for 5 minutes at 37 °C with shaking at 500 rpm. Absorbance was measured at 600 nm using a TECAN Infinite M200 PRO microplate reader (Tecan Trading AG, Switzerland), and cholesterol concentration in LP was determined from the standard calibration curve. Total lipid concentration (mg/ml) was calculated using the following formula:

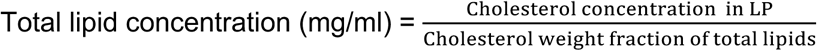

### 2.7. Primary MPN and healthy donor whole blood samples

Peripheral blood (PB) samples from patients diagnosed with MPN (n=5) were obtained from the Department of Hematology, Oncology, Hemostaseology and Stem Cell Transplantation at RWTH Aachen University Hospital following written informed consent (approved by the local ethics committee: EK 127/12). The detailed list of patients’ disease subtype, mutation profile, variant allele frequency (VAF), and treatment regimen is provided in **Table S2**. Healthy donor (HD) blood samples (n=4) were obtained from the Department of Transfusion Medicine at RWTH Aachen University Hospital, following written informed consent (approved by the local ethics committee: EK099/14). Blood samples (8 ml) were collected into EDTA tubes, stored at 4°C, and used for uptake studies within 24 hours of collection.

### 2.8. Liposome Uptake by myeloid cells in human whole blood

To evaluate the translational relevance of our findings, the uptake of LP by human whole blood was assessed. LP were prepared as described in section 2.4, incorporating 0.2 mol% DiD dye into the organic phase before mixing. For each sample, the blood collection tube was gently inverted several times to ensure homogeneity. Subsequently, 100 µl of whole blood was transferred to EDTA-containing tubes and incubated with LP at final lipid concentrations of either 250 μM or 500 μM for 1 hour in a CO_2_ incubator at 37 °C and 5 % CO_2_. Three different LP formulations (S-LP, M-LP, L-LP) were tested per donor. Following incubation, red blood cells (RBC) were lysed by adding 900 µl of red blood cell lysis solution (Erylysis buffer, Morphisto GmbH, Germany) and incubating the samples for 5 minutes at room temperature. The suspension was then transferred to FACS tubes containing 4 ml PBS to neutralize the lysis buffer. Samples were then centrifuged at 400 g for 5 minutes, and the supernatant was carefully aspirated to remove free LP and lysed cell debris. The cell pellet was washed twice with 0.5 ml FACS buffer (2% FBS, 0.5 mM EDTA in PBS), followed by centrifugation at 400 g for 5 minutes and final resuspension in 30 µl FACS buffer.

Samples were incubated with antibodies for 30 minutes in the dark at 4 °C, then washed twice and resuspended in FACS buffer. The detailed lists of staining panels and antibody dilutions are provided in **Table S3** and **Table S4**, respectively. All samples were analyzed using a BD Canto II flow cytometer (BD Biosciences, USA). Single-stain and fluorescence-minus-one (FMO) controls were included. Data were analyzed using FlowJo^™^ v10 software.

### 2.9. *JAK2*^V617F^-driven MPN murine model

All animal experiments were approved by Landesamt für Natur, Umwelt und Verbraucherschutz (LANUV of North Rhine-Westphalia (NRW, Germany), TVA 2024-409, and conducted in accordance with Directive 2010/63/EU of the European Parliament for animal experimentation.

The transgenic *JAK2*^V617F^-driven MPN murine model was established as previously described ^46^. C57BL/6 *JAK2*^V617F^ flip-flop (FF1) and C57BL/6 *JAK2* WT mice were used as BM donor mice, and C57BL/6 animals were used as recipient mice^26^. BM mononuclear cells (MNC) from donor mice were isolated after cervical dislocation under isoflurane anesthesia. Recipient mice were conditioned prior to BM transplantation with 7 Gy irradiation. Within 24 h post-irradiation, recipient mice were transplanted with 1 × 10^6^ *JAK2*^V617F^ and 1 × 10^6^ *JAK2* WT donor cells resuspended in 1x PBS (Gibco, Thermo Fisher Scientific, Germany) via tail vein injection.

To prevent infection during the post-irradiation/engraftment phase, all transplanted mice received 100 µg/ml cotrimoxazole (Cotrim K-ratiopharm^®^, Ratiopharm GmbH, Germany) in their drinking water for two weeks post-transplantation. Starting 4 weeks after transplantation, disease progression was monitored by tail-vein PB analysis every 2 weeks until week 8 post-transplantation. Blood parameters were analyzed with the Element HT5 Hematology Analyzer (Heska, USA), while flow cytometry (BD Canto II flow cytometer, BD Biosciences, USA) was used to monitor the expansion of green fluorescent protein-positive (GFP^+^) *JAK2*^V617F^ malignant clones.

At week 8 post-transplantation, a live nanoparticle biodistribution study was conducted, followed by euthanasia of the animals by cervical dislocation under isoflurane anesthesia. Organ nanoparticle accumulation was evaluated using hybrid FLT/CT imaging, while nanoparticle cellular uptake in BM and spleen-derived hematopoietic cells was analyzed by flow cytometry (LSR Fortessa Cell analyzer, BD Biosciences, USA).

### 2.10. *In vivo* biodistribution of LP formulations in a *JAK2*^V617F^ MPN murine model

Mice were randomly divided into four groups (n = 6 mice per group): free Cy7, S-LP, M-LP, and L- LP. Seven days before the start of the experiments, mice were switched to a chlorophyll-free diet to minimize background signal during optical imaging.

For *in vivo* imaging, a hybrid fluorescence / micro-computed tomography (FLT/CT) system (U-CT OI, MILabs B.V., Netherlands) was used to assess the biodistribution of Cy7-labeled LP at 0.25, 1, 4, and 24 hours post-injection. Before imaging, mice were anesthetized with 5% isoflurane (Forene^®^, Abbott, Germany) using a dedicated vaporizer and placed on a heated pad to maintain body temperature.

LP formulations were administered intravenously (i.v.) via the tail vein with a lipid concentration of 10 mM and an injection volume of 5 ml/kg, using a sterile catheter consisting of a 30 G cannula (B. Braun, Germany) connected to polyethylene tubing (inner diameter 0.28 mm, outer diameter 0.61 mm, wall thickness 0.165 mm; Hartenstein, Germany). Before imaging, ophthalmic ointment (Bepanthen^®^, Bayer Vital GmbH, Germany) was applied to prevent eye drying. Anesthetized mice were positioned in the animal holder between two acrylic plates and aligned between the FLT laser and the cooled CCD camera, with anesthesia maintained at 2 % through the imaging process.

Fluorescence scans were acquired using excitation and emission wavelengths of 730 and 775 nm, respectively, with approximately 130 scan points. Following FLT acquisition, the holder was automatically transferred to the CT module for anatomical imaging. CT scans were performed using 480 projections (1944 × 1536 pixels) over a full rotation in step-and-shoot mode at 55 kV, 0.17 mA, and an exposure time of 75 ms. This process was repeated at each time point to visualize fluorescence accumulation in major organs (liver, heart, lungs, and kidneys) and in target organs (BM and spleen). After the final imaging session, under anesthesia, blood was collected via retro-orbital sampling, followed by euthanasia by cervical dislocation. Finally, organs were harvested and processed for further *ex vivo* analysis.

The acquired 3D CT images were reconstructed at an isotropic voxel size of 80 μm using a Feldkamp filtered back-projection algorithm. Shape, scattering maps, and absorption maps were generated automatically and used for 3D fluorescence reconstruction with an established protocol^47^.

Image reconstruction and analysis were performed using Imalytics Preclinical 2.0 (Gremse-IT GmbH, Germany). 3D organ segmentation was performed on CT images using interactive tools in Imalytics Preclinical following previously established protocols^48,49^. BM, spleen, liver, heart, lungs, and kidneys were segmented for all animals at each time point. The total fluorescence signal at 0.25 h post-injection was used to calculate nanoparticle concentration (% injected dose per gram; %ID/g), from which the area under the curve (AUC) was determined and used to compare organ cumulative distribution profiles^41,50^. Means and standard deviations were calculated from n=6 mice per group. Results are presented as %ID/g as well as fold change in AUC relative to the free Cy7 group.

### 2.11. *Ex vivo* analysis of nanoparticle accumulation in BM and spleen by flow cytometry

BM cells were flushed from the femur and tibia with FACS buffer (2% FCS, 0.5 mM EDTA in 1x PBS) and mechanically dissociated by pipetting. Spleen tissue was mechanically dissociated by gently pressing tissue sections through a 70 µm cell strainer (EASYstrainer^™^, Greiner Bio-One GmbH, Germany) in FACS buffer. Cell suspensions were centrifuged at 400 g for 5 min. The cell pellet was resuspended in 1 ml of red blood cell lysis solution (Erylysis buffer, Morphisto GmbH, Germany) and incubated for 2 minutes at room temperature, followed by the addition of 5ml of 1x PBS (Gibco, Thermo Fisher Scientific, Germany) and centrifugation at 400 g for 5 min. Cell pellets were resuspended, passed through a 40 µm cell strainer (EASYstrainer^™^, Greiner Bio-One GmbH, Germany), and counted. For each FACS staining panel, 1 × 10^6^ cells were used. Staining was performed for 30 minutes at 4° C, followed by a washing step with 500 µl of FACS buffer. The detailed lists of staining panels and antibody dilutions are provided in **Table S5, Table S6,** and **Table S7**, respectively. Cells were resuspended in 350 µl FACS buffer and analyzed by flow cytometry (LSR Fortessa Cell analyzer, BD Biosciences, USA). Data were analyzed using FlowJo^™^ v10 software.

### 2.12. Histological evaluation of BM and spleen sections

The femur and spleen were harvested following euthanasia and immediately fixed in 4% paraformaldehyde (PFA). Bone samples were subsequently decalcified in 10% EDTA solution (Morphisto GmbH, Germany). Samples were submitted to the Immunohistochemistry Facility of the Interdisciplinary Center for Clinical Research (IZKF) within the Faculty of Medicine at RWTH Aachen University for routine processing, paraffin embedding, sectioning, and hematoxylin and eosin (H&E) staining. For H&E staining, the paraffin-embedded tissues were sectioned at 5 μm using a SLIDE4003E microtome (pfm Medical, Germany) and stained with H&E according to standard histological procedures. Briefly, slides containing tissue sections were deparaffinized and stained using an automated slide-staining station (ThermoFisher Scientific, Germany). The slides were stained with hematoxylin for 10 min, followed by rinsing in warm water for 10 min. The sections were then counterstained with 0.3 % eosin for 5 min, rinsed again with tap water, and then successively dehydrated in 70%, 96%, and 100% ethanol. Finally, the slides were treated with xylene and sealed with glass coverslips using Vitro-Clud® (Langenbrink GmbH, Germany).

Image acquisition was performed using the EVOSTM M7000 Imaging System (ThermoFisher Scientific, Germany). To quantify megakaryocytes, cells were counted in BM and spleen sections from three mice. Three fields of view were analyzed for each mouse, totaling nine fields per tissue. The megakaryocyte counts were then compared to the corresponding average values from healthy C57BL/6 mice.

### 2.13. Statistical analysis

All data were expressed as the mean ± standard deviation (SD) from three individual trials, unless otherwise indicated. Statistical analyses were performed using GraphPad Prism version 10.6.1 (GraphPad Software, USA). Data were analyzed by one-way ANOVA followed by Tukey’s test for multiple comparisons. Statistical significance was defined as *P < 0.05, **P < 0.01, and ***P < 0.001.

## 3. Results

### 3.1. An optimized continuous flow milli-fluidic mixing system produces liposomes with predictable physicochemical characteristics

To enable scalable LP production, a milli-fluidic system was employed^44,45^. The system, previously optimized for egg phosphatidylcholine (EPC) as the main lipid, was adapted for the production of liposomes with DPPC. As DPPC has a high transition phase temperature in comparison to EPC, the effect of operating temperature was evaluated at both 25 and 70 °C (below and above the transition temperature, respectively) to establish suitable formulation conditions.

LP composed of DPPC, cholesterol, and mPEG-(2K)-DSPE was prepared at low and intermediate lipid concentrations of 10 and 80 mM, respectively, and under defined FRR and TFR conditions. At low lipid concentration (10 mM), operating temperature had minimal impact on particle size and PDI, yielding LP of about 70 nm across the tested conditions (**Fig. S1A**). In contrast, at 80 mM, temperature had a significant effect on LP size, with LP prepared at 25°C exhibiting a larger size (118 nm ± 26.5) than LP prepared at 70 °C (76.4 nm ± 1.6) (**Fig. S1A**). This temperature dependence persisted even at high TFR (**Fig. S1B**), highlighting the importance of temperature control when formulating LP with a high transition temperature lipid, particularly at elevated lipid concentrations. Following the preliminary screening that identified 70°C as the system’s working temperature, a DoE approach was used to systematically evaluate the combined effects of three manufacturing parameters of interest, namely lipid concentration (10–150 mM), FRR, and TFR on LP size and PDI. This range encompassed both commonly used lipid concentrations (5–25 mM) and higher concentrations relevant to improving the loading efficiency of therapeutic agents ^41,44,45,51^.

Response surface modeling was used to generate contour and 3D surface plots (**Fig. 1A,B**). Within this space, LP size ranged from approximately 66 nm to 760 nm, while PDI remained low (equal to or lower than 0.2) across conditions. Increasing FRR consistently reduced LP size across the investigated lipid concentrations and TFR range, suggesting that improved mixing efficiency limits particle growth, in agreement with previous studies^41–45,52,53^. Conversely, increasing lipid concentration resulted in larger LP and higher PDI, reflecting enhanced bilayer lipid inclusion at higher lipid concentrations^44,45^. TFR showed no significant effect on particle size within the explored design space. Based on this analysis, it was apparent that lipid concentrations below 50 mM enabled the formation of liposomes up to 200 nm while maintaining low PDI (<0.2). At higher concentrations, PDI increased rapidly, particularly at lower FRR, reducing control over size. Therefore, 50 mM was selected as a practical compromise to maintain both size tunability and acceptable dispersity.

**Figure 1.**
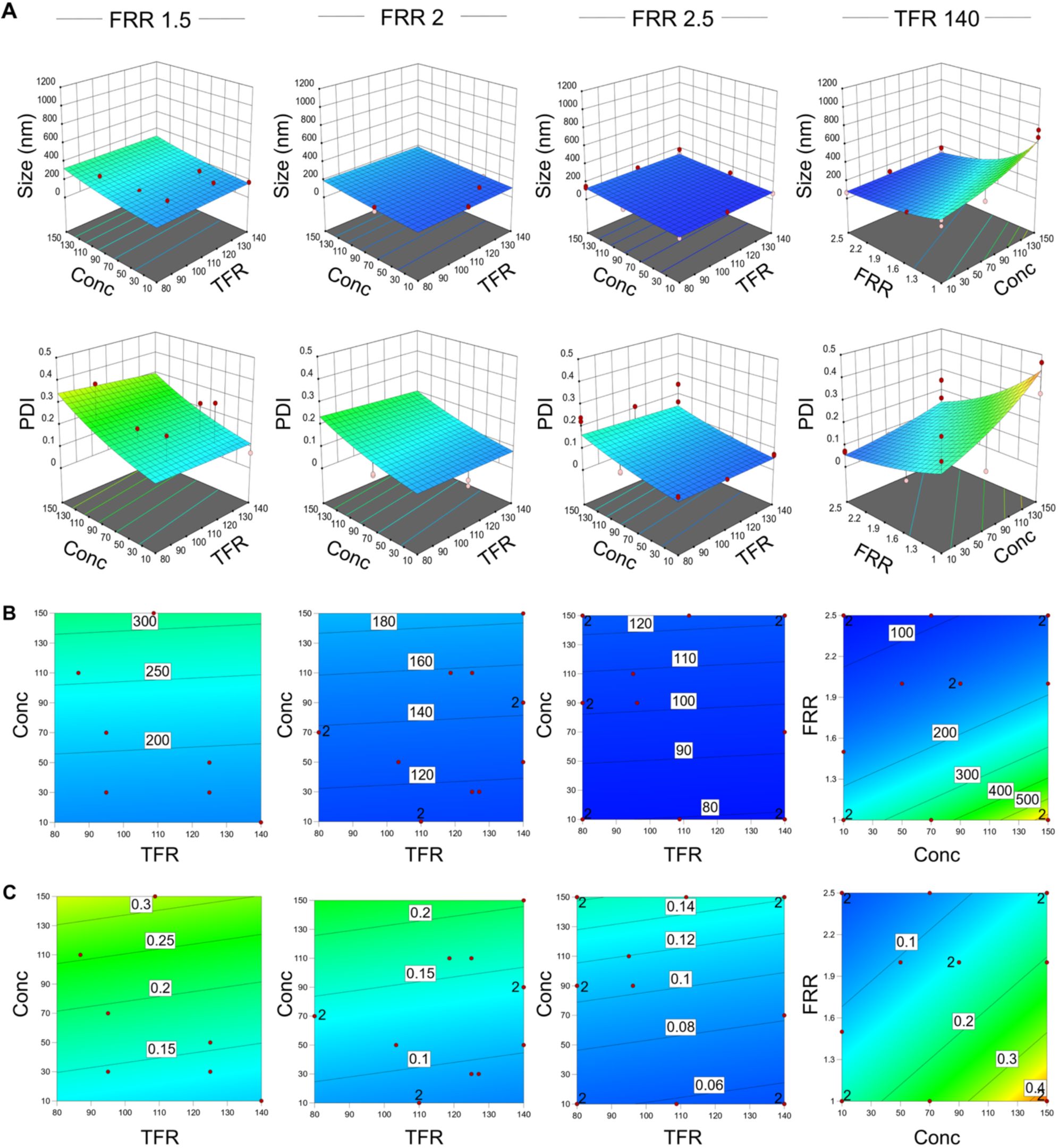
DoE surface and contour plots identify lipid concentration and FRR as critical parameters that impact LP size and PDI. (**A**) 3D-surface plots demonstrating the effect of composition parameter (lipid concentration) and process parameters (FRR and TFR) on the size and dispersity of the prepared LP. (**B-C**) 2D contour plots demonstrating the effect of composition and process parameters on LP (**B**) size and (**C**) PDI. These plots are color-coded according to response magnitude, with blue indicating lower values and yellow to red indicating higher values. The red dots represent the actual experimental conditions used to fit the DoE model. The tested ranges for the DoE were as follows: lipid concentration: 10– 150 mM; FRR: 1–2.5; and TFR: 40–140 ml/min. Detailed experimental conditions are summarized in Table S1.

The system was also evaluated regarding its compatibility with formulation additives relevant to drug loading applications. Many drugs used in the treatment of MPN and other hematological malignancies are hydrophobic or poorly soluble, making passive encapsulation (i.e., liposome formation in the presence of dissolved drug) inefficient and often resulting in low drug loading^25,41,54^. As a more efficient alternative, active encapsulation (i.e., drug encapsulation after liposome formation, using a trans bilayer gradient) is often pursued, but requires additives. To determine whether such additives affect solution viscosity, mixing behavior, and particle formation, representative dispersants, including glucose, mannitol, and ammonium sulfate, were incorporated during formulation. Importantly, their incorporation did not alter LP size (FRR 2; 103 ± 16 nm, and FRR 2.5; 71 ± 8 nm) or PDI (FRR 2; 0.16 ± 0.04, and FRR 2.5; 0.08 ± 0.03), indicating that the platform is compatible with active loading strategies and maintains particle integrity under modified formulation conditions (**Fig. S2A**).

After confirming the system modularity and defining the design space, three operating conditions were selected at a fixed lipid concentration of 50 mM and TFR of 140 ml/min. Specifically, low, moderate, and high FRR values of 1.5, 2.0, and 2.5 were selected to reproducibly generate large (L- LP; 169.1 ± 21.8 nm), medium (M-LP; 99.8 ± 5.7 nm), and small (S-LP; 61.9 ± 2.5 nm) formulations, respectively (**Fig. 2A** and **Fig. S2B**), which were then used for subsequent biological evaluation.

**Figure 2.**
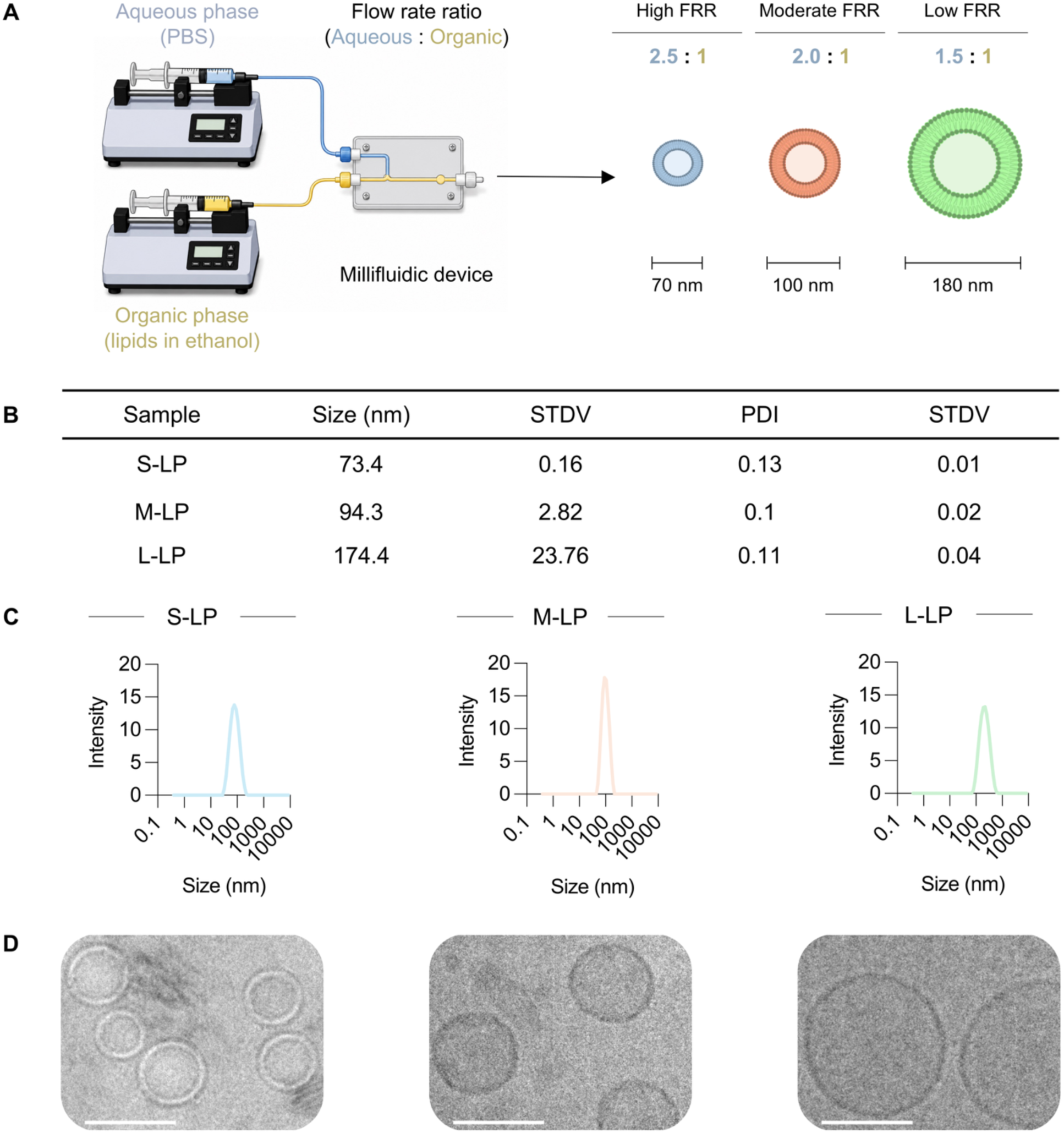
Production of Cy7-labelled LP with distinct physicochemical parameter differences, prepared via a milli- fluidic mixing device. **(A)** Schematic representation of LP preparation. LP were produced using a milli-fluidic mixing system by adjusting the FRR to obtain LP of varying sizes. **(B)** Table summarizing the mean size and PDI of the different LP formulations used in this study. LP were prepared at three defined sizes, small (S-LP), medium (M-LP), and large (L- LP). **(C)** Representative DLS charts confirming narrow size distribution across formulations. **(D)** Representative Cryo-TEM images confirm particle sizes obtained by DLS and show spherical bilayer vesicles surrounding an aqueous core, consistent with the expected morphology of liposomes. Scale bar: 100 nm.

Upon defining formulation conditions, LP with three different sizes were produced on a large scale and fluorescently labeled with a Cy7-conjugated lipid for follow-up biological evaluation (**Fig. 2A**). Cy7-labelled LP displayed hydrodynamic diameters and PDI values comparable to those of the unlabeled formulations (**Fig. 2B,C**). The morphology of LP was visualized by cryo-TEM, which revealed discrete bilayer structures enclosing an aqueous core (**Fig. 2D**), consistent with the morphology previously reported for PEGylated liposomal formulations^5,19,41,55^. The TEM images also confirmed the particle sizes determined by DLS measurements (**Fig. 2C,D**).

To enable complementary *in vitro* cellular uptake studies, LP were labeled with a DiD dye at a similar molar fraction. Similar to Cy7 incorporation, inclusion of the fluorescent lipid did not significantly affect particle size or distribution across all size classes (**Fig. S3A,B**). These findings are consistent with previous reports demonstrating that low molar fractions of fluorescent lipid dyes do not significantly impact LP size or uniformity^18–20,41^.

### 3.2. Disease state alters liposome uptake by myeloid cells in human whole blood

To assess LP uptake by human myeloid cells, LP of different size classes (S-LP = 60 nm, M-LP = 100nm, and L-LP = 180 nm) were incubated with whole blood from healthy donors (HD) or MPN patients. Uptake was quantified by flow cytometry in monocytes (CD14⁺) and granulocytes (CD66b⁺) after gating on single CD45⁺ leukocytes (**Fig. S4**).

As expected, uptake increased in a concentration-dependent manner (**Fig. 3A** and **Fig. S5**). Across all LP formulations, monocytes showed higher liposome uptake than granulocytes in both HD (**Fig. 3B,D**) and MPN patients (**Fig. 3C,E**). In HD, monocyte uptake (S-LP = 58.3 ± 15.0, M-LP = 51.5 ± 20.5, and L-LP = 25.0 ± 8.8) was approximately seven-fold higher than granulocyte uptake (S-LP = 7.5 ± 2.9, M-LP = 8.5 ± 4.3, and L-LP = 3.5 ± 1.9). In MPN patients, monocyte uptake (S-LP = 20.4 ± 11.4, M-LP = 17.12 ± 7.9, and L-LP = 10.42 ± 5.8) was approximately four-fold higher than granulocyte uptake (S-LP = 2.4 ± 1.4, M-LP = 2.5 ± 1.5, and L-LP = 1.6 ± 0.35). These findings are consistent with previous reports demonstrating higher LP affinity to human monocytes than to neutrophils in *ex vivo* whole blood^56–58^. Notably, overall uptake by myeloid cells in MPN patients (**Fig. 3C,E**) was consistently two- to three-fold lower than in HD (**Fig. 3B,D**). This reduction suggests that nanoparticle uptake is influenced by disease-associated changes in immune cell composition and function. The latter is often driven by elevated levels of pro-inflammatory cytokines, such as tumor necrosis factor alpha, which are known to alter receptor expression levels and impair the phagocytic capacity of immune cells^59–62^.

**Figure 3.**
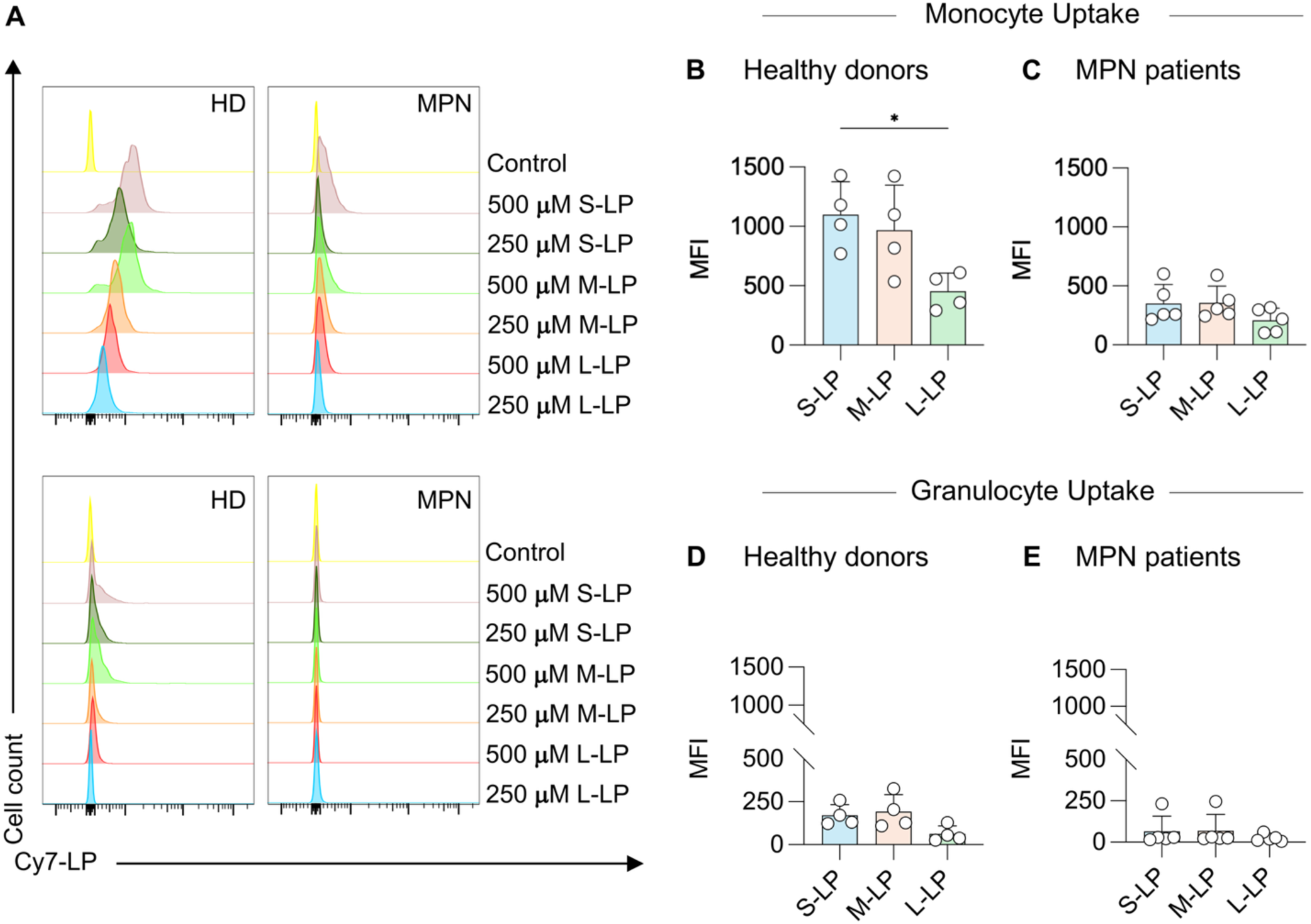
Uptake of different-sized LP by myeloid cell populations in whole blood of HD or MPN patients. Flow cytometry was used to quantify LP uptake by myeloid cells following 1 h incubation with 500 μM of the different LP formulations. (**A**) Representative flow cytometry histograms demonstrating LP uptake at two concentrations by monocytes (top row) and granulocytes (bottom row) in human whole blood. (**B-C**) LP uptake by monocytes from (**B**) HD and (**C**) MPN patients, expressed as mean fluorescence intensity (MFI) after correcting for background fluorescence by subtracting the signal measured from unstained controls. (**D-E**) LP uptake by granulocytes from (**D**) HD and (**E**) MPN patients. Across all LP formulations, monocytes exhibited higher LP uptake than granulocytes in both HD and MPN patients. Results are presented as mean ± SD (n=4). P values are indicated as * < 0.05, ** < 0.01, *** < 0.001, and **** < 0.0001.

Together, these findings underscore the importance of using pathologically relevant models to evaluate formulation performance, thereby motivating a subsequent investigation of size-dependent LP behavior *in vivo* using an MPN murine model.

### 3.3. *JAK2*^V617F^-driven MPN murine model recapitulates hallmarks of polycythemia vera

To evaluate the effect of LP size on targeting myeloid cells in hematological malignancies, we utilized an MPN mouse model. MPN phenotypic variations are largely determined by the relative expression of mutant and wild-type *JAK2* gene^26^. Elevated mutant *JAK2* signaling drives the expansion of erythroid and myeloid cell lineages. Because approximately 95% of PV patients harbor the *JAK2*^V617F^ mutation, we employed a *JAK2*^V617F^-driven PV-like murine disease model for subsequent biodistribution studies^26^.

For this, lethally irradiated mice were transplanted with a mixture of BM mononuclear cells (BMMNC) from *JAK2*^V617F^ and wild-type cells (1:1 ratio) via tail vein injection, respectively. Disease progression was monitored over 8 weeks, with PB collected every 2 weeks from week 4 to week 8 post-transplantation (**Fig. 4A**). Complete blood counts from PB were performed using an Element HT5 Hematology Analyzer (**Fig. 4B-D** and **Fig. S6A-E**).

**Figure 4.**
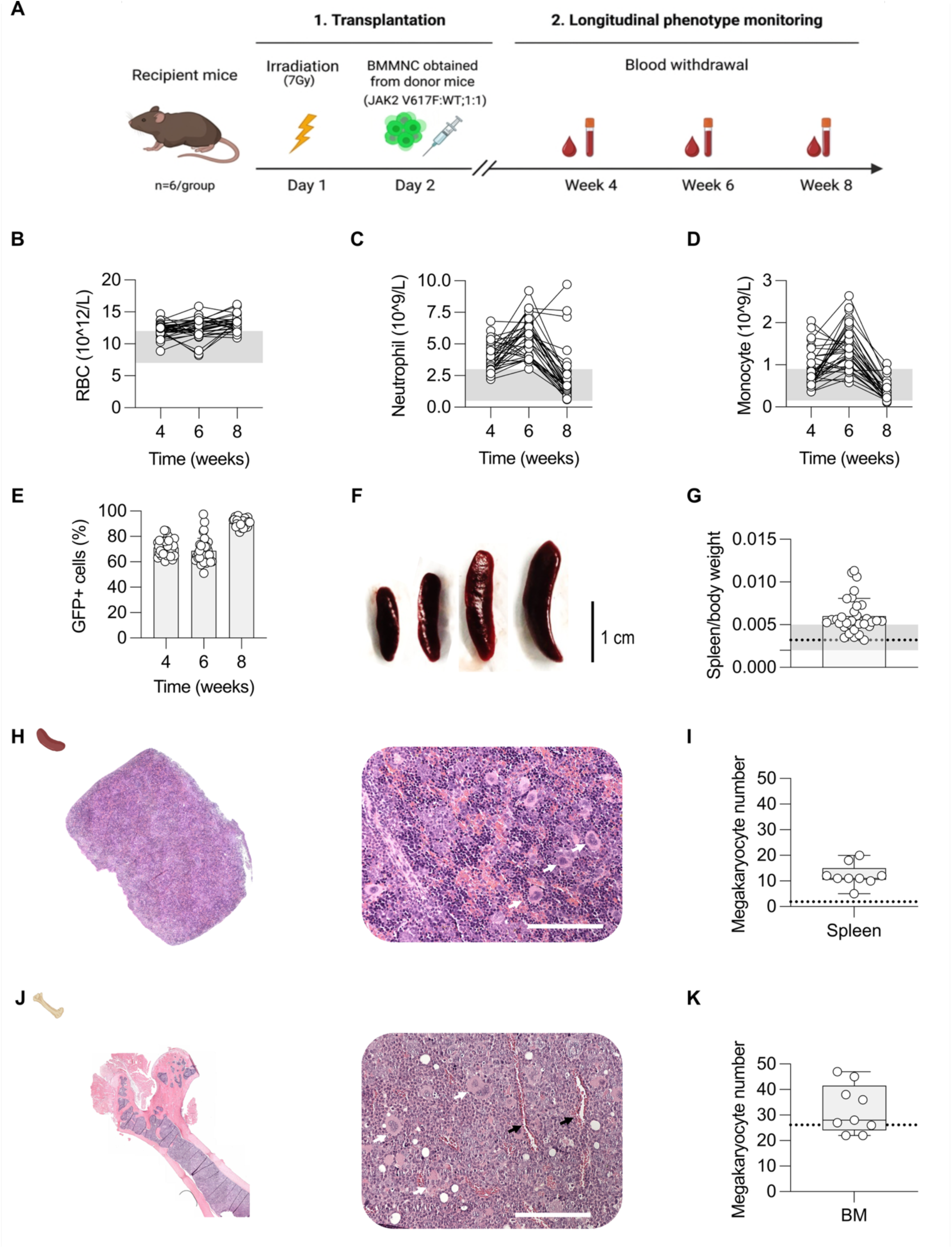
Hematological and histological analyses confirm the development of a PV-like phenotype in a *JAK2*^V617F^ MPN mouse model. (**A**) Schematic overview of the experimental design for malignant cell transplantation and longitudinal disease monitoring. Mice were lethally irradiated with 7 Gy and transplanted the following day with a 1:1 mixture of *JAK2*^V617F^ and wild-type BMMNC via tail vein injection. Complete blood counts were measured from PB collected at weeks 4, 6, and 8 post-transplantation to monitor hematologic changes associated with disease progression. Measured parameters included (**B**) RBC, (**C**) neutrophil, and (**D**) monocyte counts, all of which were elevated compared to the normal range (indicated by the grey area), demonstrating successful disease development. (**E**) Percentage of GFP⁺ donor-derived cells in the PB of transplanted mice, which increased and remained high over 8-weeks, confirming efficient engraftment and clonal expansion. (**F**) Representative images of spleens from four *JAK2*^V617F^ MPN mice demonstrating varying degrees of spleen enlargement. (**H,J**) Representative H&E stained sections of (**H**) spleen and (**J**) BM showing pronounced megakaryopoiesis (white arrow). (**I,K**) Quantification of megakaryocyte numbers per field of view in (**I**) spleen and (**K**) femoral BM sections (n=3 mice, 3 fields/mouse) showed an increase in the number of megakaryocytes in *JAK2*^V617F^ mice compared to healthy C57BL/6 mice (dotted line). Scale bar: 150 μm.

Analysis of PB revealed elevated cell counts of RBC (**Fig. 4B**), neutrophil (**Fig. 4C**), and monocyte (**Fig. 4D**) that exceeded normal reference ranges and remained elevated throughout the monitoring period, recapitulating the expected PV phenotype. In addition, engraftment efficiency was assessed by quantifying GFP^+^ donor-derived cells in PB. GFP^+^ cells accounted for 71.4% of circulating cells at week 4 and increased to 90.9% at week 8, confirming robust and stable engraftment (**Fig. 4E**). The distribution of GFP^+^ malignant cells was further assessed in spleen and BM, where engraftment efficiencies of 62.7% and 89.8%, respectively, were observed at the experiment endpoint (**Fig. S7**). Following completion of biodistribution studies at week 8, mice were euthanized, and spleens were harvested to evaluate splenomegaly due to extramedullary hematopoiesis. Marked splenomegaly was observed in the majority of animals (**Fig. 4F**), with spleen-to-body weight ratio (spleen index) consistently exceeding 0.005 (average index for healthy C57BL/6 mice, **Fig. 4G**).

Histological examination further confirmed disease progression. Spleen sections revealed marked splenomegaly with disruption of normal spleen architecture accompanied by increased extramedullary hematopoiesis and megakaryopoiesis (**Fig. 4H**). Quantification of megakaryocytes demonstrated a significant increase in their numbers in the spleen of *JAK2*^V617F^ mice compared to healthy C57BL/6 mice (**Fig. 4I**). BM sections similarly displayed enhanced megakaryopoiesis (**Fig. 4J**). Likewise, quantitative analysis confirmed the increase in megakaryocyte numbers in BM (**Fig. 4K**).

Together, these findings demonstrate the successful establishment of a PV-like MPN phenotype that recapitulates human disease and support the suitability of this model for biodistribution studies under clinically relevant inflammatory conditions.

### 3.4. Liposome size determines accumulation in BM and spleen

The BM is the primary site of disease initiation and progression in MPN^1^, while the spleen serves as a secondary hematopoietic organ that becomes increasingly involved at advanced stages, with extramedullary hematopoiesis contributing to splenomegaly^1,63–65^. Given the distinct vascular and structural characteristics of the spleen and BM^1,13,15^, we assessed the size-dependent accumulation of liposomes in these organs using a *JAK2*^V617F^ MPN model.

Cy7-labeled LP were administered i.v. (10 mM lipid concentration, 5 ml/kg body weight), and whole-body biodistribution was monitored by longitudinal 3D FLT/CT imaging at 0.25, 1, 4, and 24 h post-injection (**Fig. 5A**). Quantitative biodistribution analysis (%ID/g) showed substantial LP accumulation in the liver, as expected, as well as in the BM and spleen (**Fig. 5B,C** and **Fig. S8**). At 4 h post-injection, LP accumulation reached 6.8 % ± 3.8 %, 5.2 % ± 2.0%, and 4.4 % ± 1.8 %ID/g in the BM for L-LP, M-LP, and S-LP, respectively, and 16.6 % ± 7.7 %, 14.3 % ± 8.1 %, and 9.4 % ± 5.18 %ID/g in the spleen (**Fig. 5B**). In both organs, LP accumulation increased with particle size, with L-LP consistently exhibiting the highest accumulation, followed by M-LP and S-LP. LP accumulation remained higher in the spleen than in the BM throughout the imaging period.

**Figure 5.**
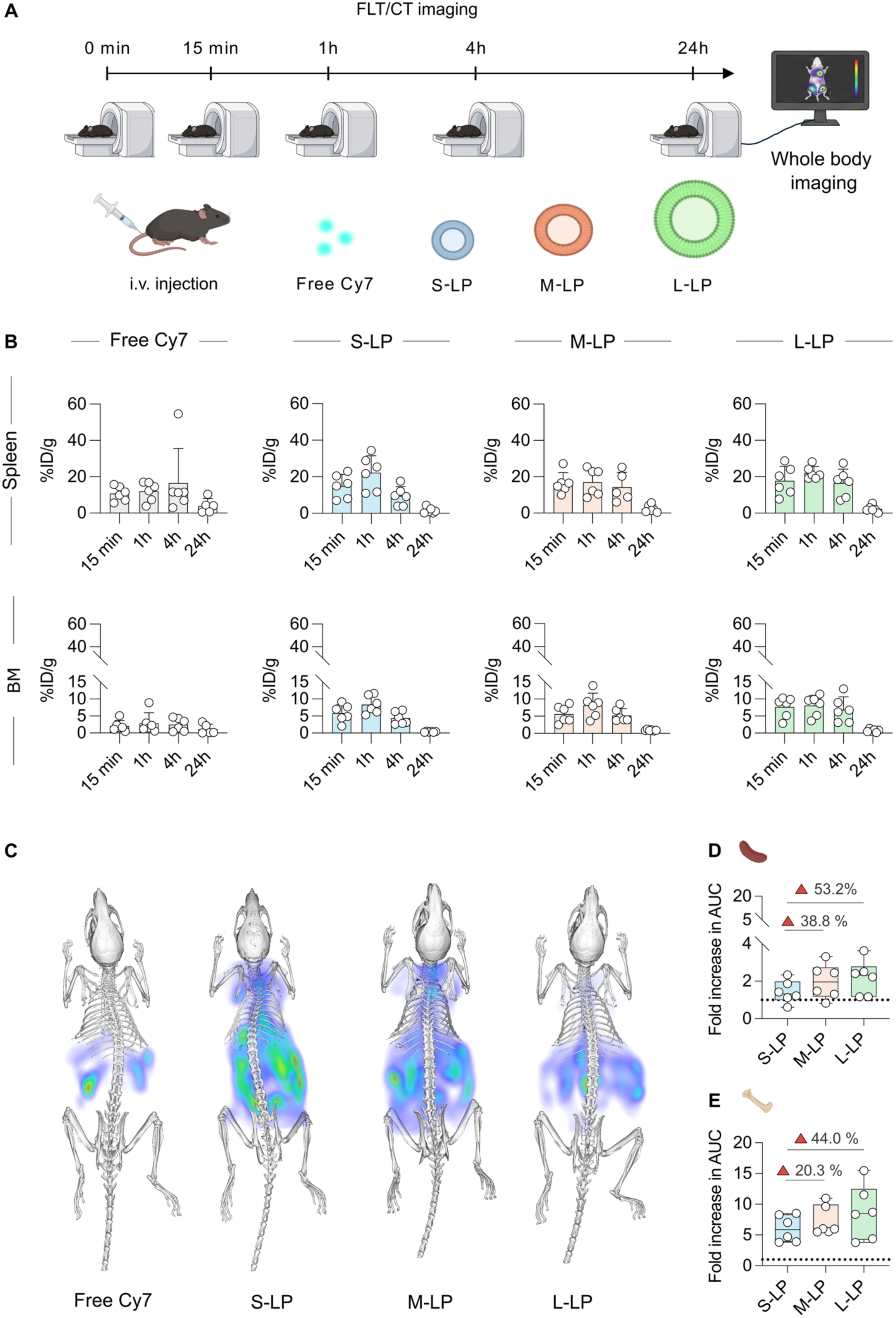
Liposome size drives delivery to hematopoietic tissues (spleen and BM) in a *JAK2*^V617F^ MPN mouse model. (**A**) Schematic overview of the study design. All LP formulations were injected i.v. through the tail vein at t = 0 min, followed by longitudinal *in vivo* FLT/CT imaging at the indicated time points. (**B**) Quantitative biodistribution of the different- sized Cy7-labeled LP, expressed as %ID/g, in the spleen and BM at 15 min, 1 h, 4 h, and 24 h post-injection, illustrating the accumulation kinetics of LP in both hematopoietic organs. (**C**) Representative FLT/CT images acquired at 24 h after i.v. administration of free Cy7, S-LP, M-LP, or L-LP, depicting the biodistribution of the different LP formulations. (**D,E**) LP accumulation in the (**D**) spleen and (**E**) BM expressed as fold increase in AUC relative to free Cy7 (dotted line), demonstrating increased cumulative accumulation with increasing LP size. Results are presented as mean ± SD; with n = 6 per group.

Since %ID/g reflects organ accumulation at individual time points, cumulative tissue exposure over the entire imaging period was assessed by calculating the AUC (%ID/g*h) for each formulation and then normalizing it to the free Cy7 control (**Fig. 5D,E**). Consistent with the %ID/g analysis, a size-dependent increase in LP accumulation was observed in both hematopoietic organs, with L-LP exhibiting the highest AUC, followed by M-LP and S-LP. Relative to S-LP, M-LP and L-LP increased the AUC by 38.8% and 53.2%, respectively, in the spleen, whereas the corresponding increases in the BM were 20.3% and 44.0%. When compared with the free model-drug dye, the relative enhancement in AUC was greater in the BM (LP = 8.7 ± 4.4, M-LP = 7.3 ± 2.4, S-LP = 6.0 ± 2.16) than in the spleen (LP = 2.2 ± 0.9, M-LP = 2.0 ± 0.9, S-LP = 1.4 ± 0.6) (**Fig. 5D,E**).

Together, these findings demonstrate that increasing LP size enhances its accumulation within both hematopoietic organs, with the greatest improvement observed in the BM compared with the free model-drug dye.

### 3.5. Increasing liposome size enhances uptake by myeloid cells in hematopoietic organs

Following organ-level nanoparticle biodistribution assessment, *ex vivo* analysis of BM, spleen, and PB was performed for evaluating nanoparticle uptake by myeloid immune cells using flow cytometry at 24 h post-injection (**Fig. 6A**). Across all tissues, LP uptake was dependent on particle size, with L-LP showing significantly higher uptake by CD11b⁺Gr1⁺ myeloid cells (monocytes and granulocytes; **Fig. 6B**), CD11b⁺Ly6G⁺ neutrophils (**Fig. 6C**), and CD11b⁺Gr1^-^Ly6G^-^ monocytes/macrophages (**Fig. 6D**).

**Figure 6.**
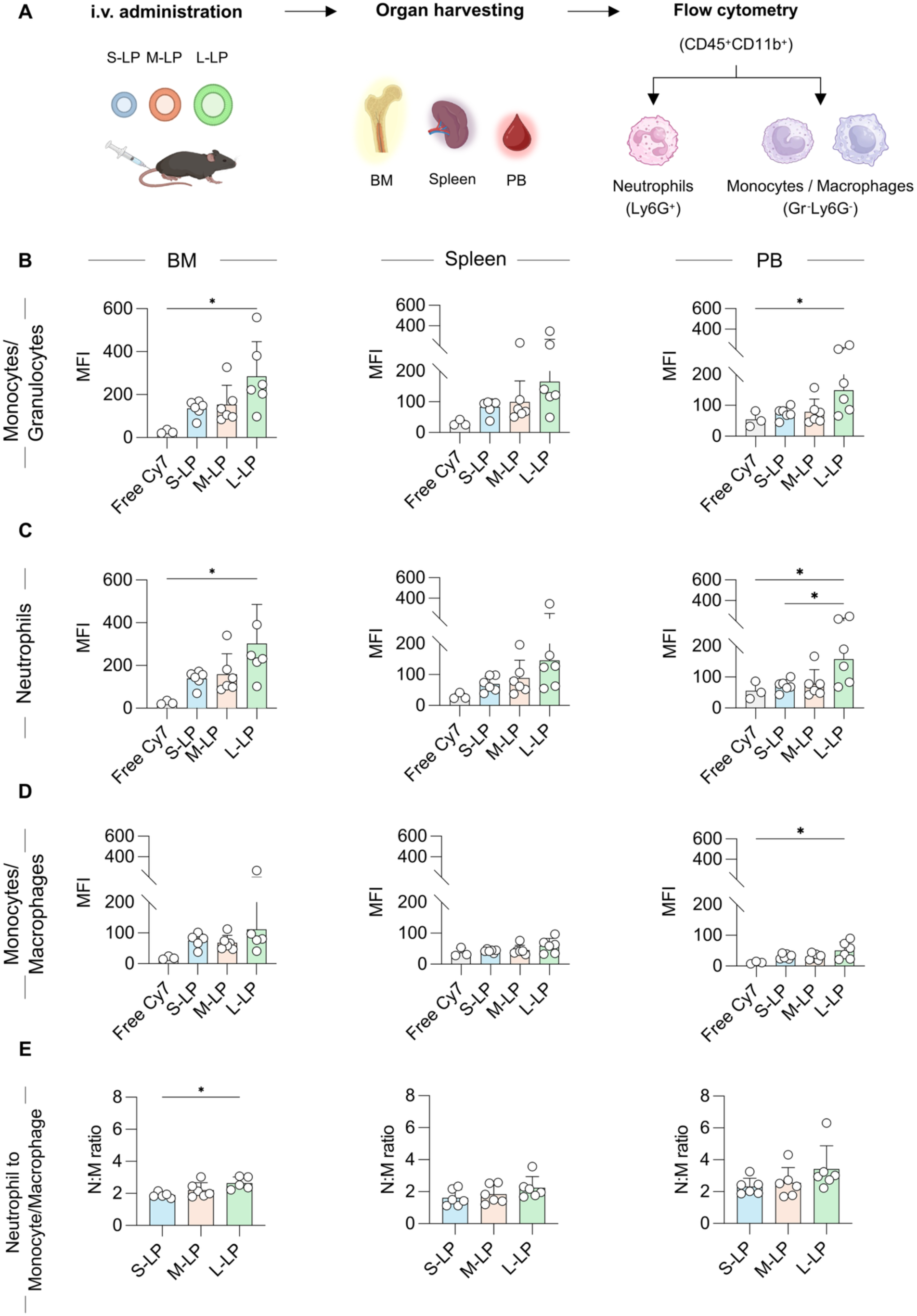
Increasing liposome size enhances uptake by myeloid cells across hematopoietic organs and peripheral blood. **(A)** Schematic overview of the experimental workflow, including organ collection, flow cytometric analysis, and the gating strategy employed to identify the different myeloid populations. LP uptake was evaluated in (**B**) myeloid cells (monocytes and granulocytes), (**C**) neutrophils, and (**D**) monocytes/macrophages, across BM, spleen, and PB. The data demonstrated that LP uptake increased with particle size, with L-LP consistently exhibiting the highest uptake across hematopoietic compartments, particularly in the BM. (**E**) Neutrophil uptake relative to monocyte and macrophage was assessed by calculating neutrophil to monocyte/macrophage (N:M) ratio. The results indicated that neutrophils consistently exhibited greater LP uptake than both monocytes and mature macrophages across all organs, with the highest ratios observed for L-LP. Results are presented as mean ± SD (n=5-6). P values are indicated as * < 0.05, ** < 0.01, *** < 0.001, and **** < 0.0001.

In the BM, L-LP showed higher uptake compared to M-LP and especially S-LP, for both monocytes/macrophages (L-LP = 112.06 ± 88.42, M-LP = 68.15 ± 24.07, S-LP = 74.80 ± 23.92) and neutrophils (L-LP = 302.5 ± 183.8, M-LP = 159.2 ± 95.4, S-LP = 138.5 ± 36.6) (**Fig. 6C,D** and **Fig. S9**). A similar trend was observed for both monocytes/macrophages and neutrophils in PB and spleen (**Fig. 6C,D**).

Across compartments, overall cellular uptake was highest in the BM, revealing a two-fold increase in comparison to spleen and PB (**Fig. 6** and **Fig. S9**). At 24 h post-injection, these differences are not trivial, as they indicate that although nanoparticles are cleared from circulation and clearance organs, they are retained to a greater extent in the BM, which is the disease hot-spot for inflammatory and malignant manifestations.

Another important comparison is the uptake in neutrophils versus monocytes/macrophages (**Fig. 6E** and **Fig. S9**). Neutrophils displayed higher uptake than both populations, with neutrophil over monocyte/macrophage (N:M) uptake ratios >1 across all organs and formulations. Although the absolute LP uptake (MFI) of both neutrophils and monocytes/macrophages was highest in the BM, the relative difference between these populations was most pronounced in the peripheral blood, particularly for L-LP administration. Specifically, the N:M uptake ratio was higher in the PB (L-LP = 3.42 ± 1.46, M-LP = 2.55 ± 0.95, S-LP = 2.32 ± 0.52), compared with the BM (L-LP = 2.64 ± 0.39, M-LP = 2.20 ± 0.46, S-LP = 1.89 ± 0.17) and spleen (L-LP = 2.24 ± 0.70, M-LP = 1.83 ± 0.58, S-LP = 1.61 ± 0.52), indicating a greater relative difference in LP uptake between neutrophils and monocyte/macrophages in these tissues (**Fig. 6E**).

Finally, uptake by both lineage-negative (Lin^⁻^) and the hematopoietic stem and progenitor-enriched cell populations Lin⁻Sca-1⁺c-Kit⁺ (LSK^+^) was minimal, as expected, given the restricted accessibility of these cells within the BM and splenic niche as well as their limited phagocytic capacity (**Fig. S10**). Taken together, these data indicate that increased LP size results in extensive uptake by myeloid cells in hematopoietic organs, with more prominent uptake by neutrophils. The high N:M uptake ratio (in all cases >1) indicates that MPN neutrophils play an important role in engaging with nanomaterials. This becomes even more significant considering their higher abundance compared to monocytes and macrophages.

## 4. Discussion

In this study, we demonstrated that nanoparticle size is a key formulation parameter that influences engagement with myeloid cells, and hematopoietic organ biodistribution in hematological malignancies. More specifically, we showed that size-dependent differences in biodistribution were linked to interactions with myeloid cells, particularly neutrophils. This is important in MPN, as these populations contribute to disease pathobiology and infiltrate key compartments, such as the BM and spleen. Hence, their capacity to internalize nanoparticles highlights their potential as biologically relevant targets and supports the translational relevance of the observed uptake patterns.

To achieve precise and reproducible control over liposome size, we employed a continuous flow manufacturing strategy using milli-fluidics. This was essential, as conventional nanoparticle preparation methods are limited in scalability and make it challenging to produce larger particles. In contrast, the milli-fluidic approach allowed us to generate liposomes across a wide size range with small PDI, thereby enabling assessment of size-dependent biological effects. Optimization of milli- fluidic parameters identified FRR and lipid concentration as the main variables influencing liposome size and dispersity. Increasing FRR resulted in smaller and more uniform liposomes, likely due to faster ethanol dilution and narrower mixing streams that limit particle fusion and growth^41,66^. In contrast, higher lipid concentrations promoted the formation of larger and more polydisperse vesicles, likely due to inefficient mixing between the organic and aqueous phases, leading to uncontrolled vesicle growth and increased fusion of bilayers^44,67^.

In *ex vivo* studies of whole-blood samples, liposome uptake was primarily mediated by monocytes, which is in agreement with prior studies in both healthy and diseased human and murine subjects^19–21,56–58,68–70^. In addition, overall uptake was reduced in MPN samples compared to HD. While chronic inflammation in MPN is known to affect immune cell function, direct comparisons of nanoparticle uptake between HD and patients remain limited^56,59–62^. Here, our data reveal the first (to the best of our knowledge) evidence that disease state can reduce nanoparticle uptake, underscoring the importance of evaluating nanomedicine performance in pathologically relevant systems.

While informative, these *ex vivo* blood assays usually capture initial early interactions between nanoparticles and circulating immune cells under static conditions. Under these conditions, particularly at short incubation times, monocyte contributions to nanoparticle uptake are likely to be overrepresented, whereas neutrophil involvement may be underestimated^69^. In contrast, the *in vivo* setting incorporates physiological processes that cannot be recapitulated *ex vivo*, including dynamic changes in circulating immune cell populations following intravenous administration, immune cell trafficking, vascular interactions, and tissue homing, all of which influence nanoparticle interactions with immune cells and distribution^57^. Supporting this, a previous murine study comparing liposome association in *ex vivo* and *in vivo* blood demonstrated that total neutrophil association was more than 50-fold higher *in vivo* after 1 h^57^. Hence, to account for these additional biological processes and better understand liposome behavior under physiologically relevant inflammatory conditions, we employed a murine *JAK2*^V617F^ MPN model that recapitulates the human PV phenotype. This model was characterized by sustained leukocytosis, erythrocytosis, neutrophilia, and monocytosis as confirmed by PB analysis and histological assessment of the spleen and BM.

Whole-body hybrid imaging revealed that liposome size strongly influences biodistribution, with a trend of larger particles preferentially accumulating in myeloid-rich organs, such as the BM and spleen. The increased splenic accumulation of larger particles is consistent with the established size-based filtration function of this organ, where particles larger than 150 nm are unable to traverse the endothelial slit of splenic sinuses and are therefore retained in the red pulp, where they are engulfed by red pulp macrophages^13,71^. This high splenic accumulation may be of clinical relevance since MPN patients frequently develop splenomegaly, particularly in advanced stages of the disease.

On the other hand, the preferential accumulation of L-LP in the BM is less intuitive, as passive transport mechanisms, such as passive diffusion through endothelial fenestrae, which typically range from 50–150 nm, are generally expected to favor smaller particles^1,16,72–74^. To better understand this observation, we performed cellular uptake analysis, which confirmed that liposome uptake increased with particle size and further revealed that larger liposomes preferentially engaged with myeloid cells (monocytes and granulocytes) across BM, spleen, and peripheral blood. This observation is consistent with previous reports indicating that myeloid cells interact with larger particles at higher extents^11,56,57,75–78^. These findings further suggest that myeloid cells contribute to the delivery of liposomes to the BM. This interpretation is supported by emerging evidence that myeloid cells can actively mediate nanoparticle transport under inflammatory conditions, a process often referred to as immune-cell “hitchhiking”^18–20,22^. While this mechanism is typically described in the context of targeting inflammatory lesions and tumors, our findings suggest that a similar process may facilitate liposome delivery to the BM. In the context of MPN, this mechanism may be supported by the chronic inflammatory state of the disease, which is characterized by enhanced granulopoiesis, monocytosis, and increased cytokine signaling. These processes promote both neutrophil turnover and clearance as well as monocyte recruitment to hematopoietic tissues^28–34^.

Senescent neutrophils are known to return to the BM for clearance ^31,32^, and during this process, may transport internalized liposomes back to this compartment. Indeed, the intrinsic marrow-homing behavior of neutrophils has recently been exploited for the delivery of polymeric nanoparticles^22^. At the same time, inflammatory recruitment of monocytes to BM may also contribute to the observed accumulation pattern^33,34^. Together, these findings suggest that myeloid cells may function as endogenous carriers that facilitate liposome transport and accumulation within the BM^31,32^.

Previous strategies aimed at targeting the BM have largely focused on monocyte populations, either through formulation design that specifically targets these cells or experimental setups that emphasize or, in some cases, bias the role of monocytes in mediating nanoparticle uptake^79,80^. The findings presented here expand the current understanding by identifying neutrophils alongside monocytes as contributors to liposome accumulation in the BM. More importantly, in disease settings characterized by enhanced granulopoiesis, neutrophils may represent a so-far underappreciated pathway for nanoparticle delivery to this compartment.

Taken together, our findings identify particle size as a determinant of nanoparticle distribution in hematologic disease through engagement with myeloid cells, and especially neutrophils. This is notable, as nanoparticle design is often guided by principles derived from solid tumors, where accumulation is largely driven by vascular permeability. Instead, our results highlight the need to incorporate disease-specific biology into the design of nanoparticle-based therapies and suggest that size-dependent engagement of myeloid cells may be leveraged to improve delivery to hematopoietic compartments in oncology and beyond.

## 5. Conclusions

This study combines scalable liposome manufacturing, process optimization, and disease-relevant biological evaluation to investigate how particle size influences nanoparticle behavior in hematologic malignancies. Using a milli-fluidic platform optimized through a DoE approach, we generated liposomes with defined size classes and evaluated liposome uptake *ex vivo* in human whole blood and *in vivo* in a murine *JAK2*^V617F^ MPN model. By linking manufacturing parameters to particle size and ultimately to cell-specific biodistribution patterns in a murine model of hematologic malignancy, we show that particle size strongly influences biodistribution through immune cell-mediated mechanisms. In particular, our results highlight neutrophils as key contributors to liposome accumulation in the BM, suggesting that neutrophil endogenous trafficking may be harnessed as a natural transport mechanism to direct nanoparticles to this site. It also expands our understanding of neutrophil targeting beyond ligand-based strategies. Together, these mechanistic insights have important implications for the rational design of nanomedicines intended for hematologic malignancies and other BM-related disorders.

## Data availability

The data are available from the authors upon request.

## Author contribution

AMS, MAST, and JMM conceptualized and supervised the study. SE, MAST, AMS, and JMM designed the experiments. SE designed and executed the mathematical modeling. SE developed and characterized the LP formulations. EMB performed Cryo-TEM imaging. SKos treated the MPN patients and provided the blood samples. SE performed the uptake studies on human whole blood samples. SE and MAST performed the flow cytometry experiments and analyzed the data on *ex vivo* blood samples. MAST developed the MPN mouse model, monitored disease progression, and performed the histological analysis. MAST, AMS, JB, SE, AN, ShK, FD, and AM conducted the animal experiments. MAST, MV, MA, and JL performed the flow cytometry experiments and analyzed the data for the animal study. SE, ShK, FD, and AMS analyzed and validated the FLT/CT scans. SE performed statistical analyses. SE drafted the manuscript. SE, MAST, and AMS prepared the figures. AMS, MAST, JMM, JB, FD, NC, SKos, TL, and FK provided funding. All authors contributed to data interpretation, manuscript revision, and approved the final version for submission.

## Acknowledgements

The authors acknowledge financial support from the German Research Foundation (DFG; Excellence Initiative RWTH JPI 2021 (AMS), Clinical Research Unit (CRU) 344 (428858786; NC, AMS, SKos), GRK 2375 Tumor-Targeted Drug Delivery (331065168; SE, JMM, FK, TL)), German José Carreras Leukemia Foundation (DJCLS 06R/2024; MAST), and the German Cancer Aid through the Center for Integrated Oncology Aachen Bonn Cologne Düsseldorf (CIO^ABCD^) at the Mildred Scheel School of Oncology (MSSO) (SDK; CIO^ABCD-MSSO^ (FD, JB, AMS, TL, SKos)). The flow cytometry measurements were supported by the Flow Cytometry Facility and Immunohistochemistry Facility, core facilities of the Interdisciplinary Center for Clinical Research (IZKF) Aachen within the Faculty of Medicine at RWTH Aachen University. The schematics were created with BioRender.com.

