## Supplementary Information for "Nanoparticle size governs engagement with myeloid cells and hitchhiking to hematopoietic organs in myeloproliferative neoplasms"

**Table S1**. Number of runs, lipid concentration, flow rate ratio (FRR), and total flow rate (TFR) determined by DoE software, along with the corresponding measured size and PDI.

| Block | Run # | Lipid concentration | FRR | TFR (ml/min) | Size (nm) | PDI |
|---|---|---|---|---|---|---|
| Block 1 | 1 | 10 | 1 | 140 | 191.2 | 0.217 |
| Block 1 | 2 | 70 | 1 | 110 | 373.3 | 0.473 |
| Block 1 | 3 | 10 | 2.5 | 80 | 72.91 | 0.086 |
| Block 1 | 4 | 10 | 2.5 | 140 | 66.04 | 0.068 |
| Block 1 | 5 | 150 | 2 | 140 | 159 | 0.154 |
| Block 1 | 6 | 10 | 1 | 140 | 250.8 | 0.316 |
| Block 1 | 7 | 10 | 2.5 | 80 | 67.46 | 0.081 |
| Block 1 | 8 | 10 | 2.5 | 140 | 67.36 | 0.073 |
| Block 1 | 9 | 150 | 1 | 80 | 584.7 | 0.311 |
| Block 1 | 10 | 50 | 2 | 103.4 | 107.9 | 0.079 |
| Block 1 | 11 | 10 | 1 | 80 | 375.5 | 0.309 |
| Block 1 | 12 | 150 | 1 | 140 | 608 | 0.467 |
| Block 1 | 13 | 110 | 2 | 118.7 | 105.6 | 0.065 |
| Block 1 | 14 | 110 | 1.5 | 86.9 | 314 | 0.409 |
| Block 1 | 15 | 150 | 2.5 | 80 | 150 | 0.241 |
| Block 1 | 16 | 150 | 2.5 | 111.6 | 142.5 | 0.211 |
| Block 2 | 17 | 150 | 1.5 | 108.8 | 228.4 | 0.305 |
| Block 2 | 18 | 70 | 2.5 | 140 | 121.7 | 0.074 |
| Block 2 | 19 | 10 | 2.5 | 108.8 | 104 | 0.06 |
| Block 2 | 20 | 150 | 1 | 80 | 577.6 | 0.418 |
| Block 2 | 21 | 150 | 2.5 | 140 | 118.8 | 0.162 |
| Block 2 | 22 | 10 | 1 | 80 | 217.5 | 0.122 |
| Block 2 | 23 | 30 | 2 | 127.1 | 143.6 | 0.058 |
| Block 2 | 24 | 150 | 1 | 140 | 759.8 | 0.332 |
| Block 2 | 25 | 150 | 2.5 | 80 | 130.5 | 0.224 |
| Block 2 | 26 | 10 | 1 | 80 | 140.6 | 0.096 |
| Block 2 | 27 | 90 | 2.5 | 96.2 | 81.65 | 0.055 |
| Block 2 | 28 | 70 | 2 | 80 | 160.5 | 0.08 |
| Block 2 | 29 | 70 | 1 | 140 | 237.1 | 0.092 |
| Block 2 | 30 | 150 | 2.5 | 140 | 178.6 | 0.251 |
| Block 2 | 31 | 10 | 1.5 | 140 | 171 | 0.071 |
| Block 3 | 32 | 70 | 1.5 | 95 | 220.4 | 0.238 |
| Block 3 | 33 | 90 | 2.5 | 80 | 90.7 | 0.07 |
| Block 3 | 34 | 90 | 2.5 | 80 | 91.15 | 0.075 |
| Block 3 | 35 | 30 | 1.5 | 95 | 249.5 | 0.256 |
| Block 3 | 36 | 90 | 2 | 140 | 92.91 | 0.069 |
| Block 3 | 37 | 50 | 2 | 140 | 112.9 | 0.069 |
| Block 3 | 38 | 10 | 2 | 110 | 133.7 | 0.019 |
| Block 3 | 39 | 10 | 2.5 | 95 | 92.55 | 0.085 |
| Block 3 | 40 | 70 | 2 | 80 | 112.2 | 0.086 |
| Block 3 | 41 | 110 | 2 | 125 | 131.1 | 0.084 |
| Block 3 | 42 | 30 | 1.5 | 125 | 212.2 | 0.316 |
| Block 3 | 43 | 30 | 2 | 125 | 105.3 | 0.066 |
| Block 3 | 44 | 50 | 1.5 | 125 | 281.4 | 0.293 |
| Block 3 | 45 | 10 | 2 | 110 | 128 | 0.045 |
| Block 3 | 46 | 90 | 2 | 140 | 106 | 0.073 |

**Table S2.** Details the disease subtype, mutation profile, variant allele frequency (VAF), and treatment regimen of MPN patients whose peripheral blood samples were analyzed in this study.

| Patient ID | MPN disease entity | Mutation profile | VAF (%) | Treatment |
|---|---|---|---|---|
| 1 | Post-PV-MF | JAK2 V617F | 72 | Ruxolitinib |
| 2 | PV | JAK2 V617F | 61 | No treatment |
| 3 | PV | JAK2 V617F | 68 | Hydroxyurea + Interferon |
| 4 | PV DD pre-PMF | JAK2 V617F | 87 | Hydroxyurea |
| 5 | PV | JAK2 V617F | 50 | No treatment |
| | | TET2 1-Bp Del (p.Lys1439AsnfsTer9); | 27 | |

**Table S3.** Summary of antibody targets, fluorophores, and suppliers used in flow cytometry analyses of human whole blood samples.

| Human FACS antibodies | | |
|---|---|---|
| **Target** | **Fluorochrome** | **Company** |
| CD45 | PB | Biolegend |
| CD14 | PE | Biolegend |
| CD66b | PE | Biolegend |

**Table S4.** Antibody targets, fluorophores, and dilutions used for identifying granulocyte and monocyte populations in human blood samples for flow cytometry–based quantification of nanoparticle uptake.

| **Human Granulocyte/Monocyte panel** | | |
|---|---|---|
| **Target** | **Fluorochrome/Channel** | **Dilution** |
| Nanoparticles | APC-Cy7 | - |
| CD45 | PB | 1 to 100 |
| CD14 | PE | 1 to 100 |
| CD66b | PE | 1 to 100 |

**Table S5**. List of the targets, fluorophores, and suppliers of mouse antibodies used for flow cytometry–based quantification of nanoparticle accumulation in ex vivo samples.

| Murine FACS antibodies | | |
|---|---|---|
| **Target** | **Fluorochrome** | **Company** |
| CD45 | PB | Biolegend |
| Gr1 | PE | Biolegend |
| CD11b | PE-Cy7 | Biolegend |
| Ly6G | APC | Biolegend |
| c-KIT | APC | Biolegend |
| Sca1 | PE-Cy7 | Biolegend |
| CD11b | PE-Cy5 | Biolegend |
| Gr1 | PE-Cy5 | Biolegend |
| B220 | PE-Cy5 | Biolegend |
| CD4 | PE-Cy5 | Biolegend |
| CD8 | PE-Cy5 | Biolegend |
| CD3 | PE-Cy5 | Biolegend |
| Ter119 | PE-Cy5 | Biolegend |

**Table S6.** Antibody targets, fluorophores, and dilutions used for identifying granulocyte and monocyte populations in ex vivo samples by flow cytometry.

| Granulocyte/Monocyte panel | | |
|---|---|---|
| **Target** | **Fluorochrome/Channel** | **Dilution** |
| Nanoparticles | APC-Cy7 | - |
| Transplanted cells | GFP | - |
| CD45 | PB | 1 to 100 |
| Gr1 | PE | 1 to 100 |
| CD11b | PE-Cy7 | 1 to 100 |
| LyG | APC-Cy7 | 1 to 100 |

**Table S7.** List the antibody targets, corresponding fluorochromes, detection channels, and antibody dilutions used for flow cytometry.

| LSK panel | | |
|---|---|---|
| **Target** | **Fluorochrome/Channel** | **Dilution** |
| Nanoparticles | APC-Cy7 | - |
| Transplanted cells | GFP | - |
| c-KIT | APC | 1 to 100 |
| Sca-1 | PE-Cy7 | 1 to 100 |
| CD11b | PE-Cy5 | 1 to 200 |
| Gr1 | PE-Cy5 | 1 to 200 |
| B220 | PE-Cy5 | 1 to 200 |
| CD4 | PE-Cy5 | 1 to 200 |
| CD8 | PE-Cy5 | 1 to 200 |
| CD3 | PE-Cy5 | 1 to 200 |
| Ter119 | PE-Cy5 | 1 to 200 |

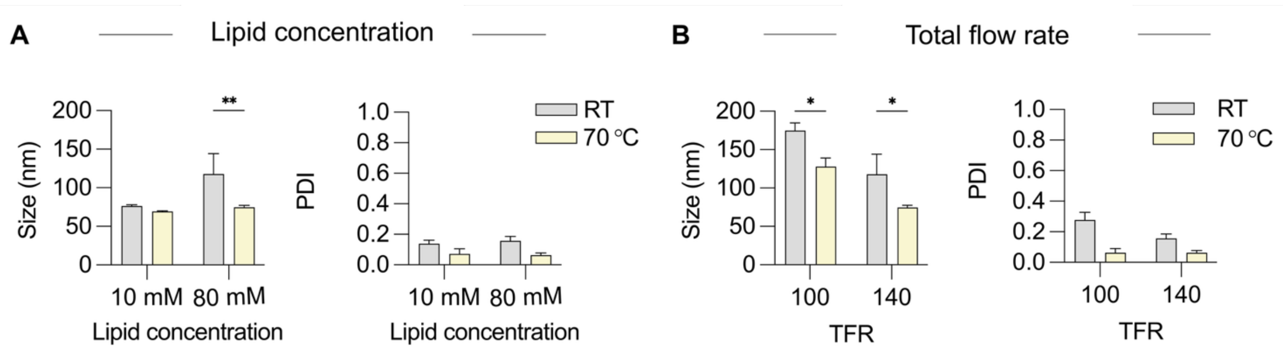


**Figure S1. Elevated temperatures (above DPPC phase transition temperature) promote the formation of smaller, more uniform liposomes.** Size and PDI of liposomes prepared with (**A**) varying lipid concentrations and (**B**) different total flow rates (TFR) and prepared either at room temperature (RT) or 70 °C. Increasing the system temperature resulted in smaller liposome sizes and lower PDI values, particularly at higher lipid concentrations. A similar trend was observed with increasing TFR from 100 to 140 mL/min. Results are expressed as mean ± SD (n = 3). P values are indicated as * < 0.05, ** < 0.01, *** < 0.001, and **** < 0.0001.

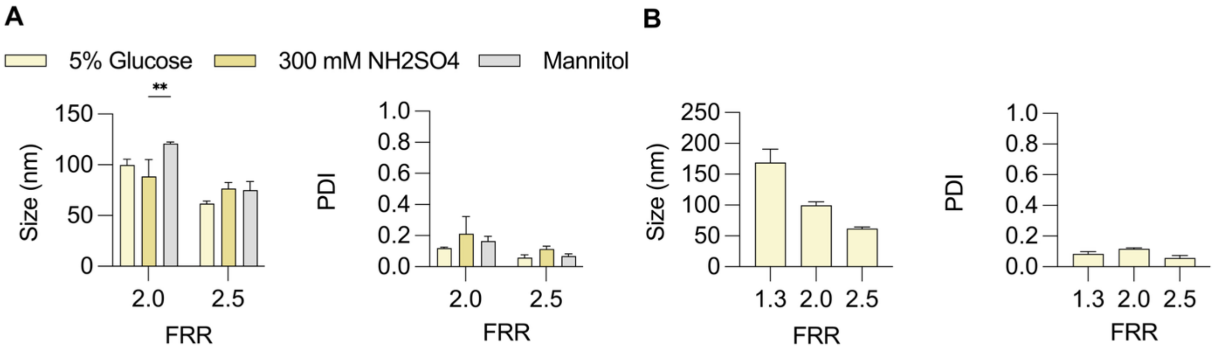


**Figure S2. Evaluation of the modularity of the CFM for drug loading.** (**A**) Effect of additives, namely 5% glucose, ammonium sulfate $(NH_4)_2SO_4$, and mannitol on LP size and PDI, confirming the modularity and compatibility of CFM for active drug loading. (**B**) The three selected FRR that were used to generate LP with target sizes of 70, 100, and 180 nm. Results are expressed as mean ± SD (n = 3).

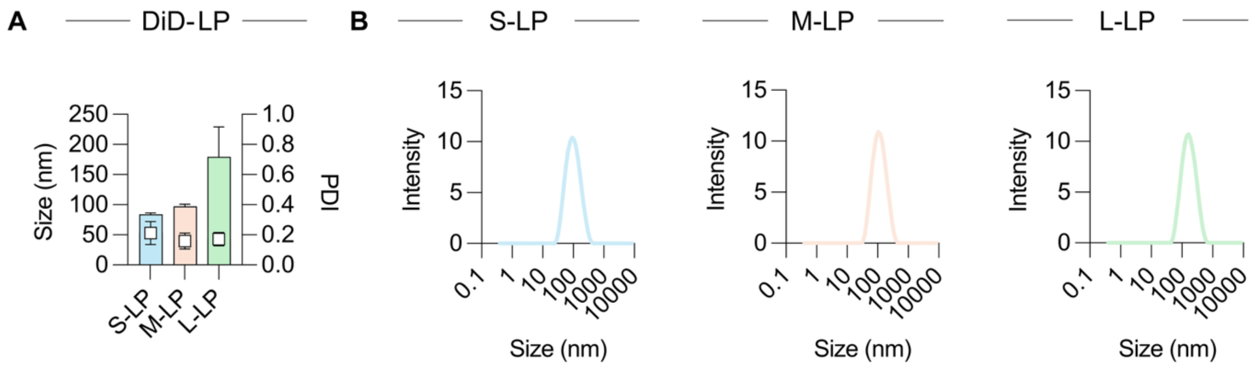


**Figure S3. Characterization of DiD-labeled LP of different sizes used for uptake studies in human whole blood.** (**A**) Size and PDI of DiD-labeled LP as measured by dynamic light scattering (DLS). (**B**) Representative DLS intensity distribution charts demonstrating narrow size distribution across formulations. Data are presented as mean ± SD (n = 3).

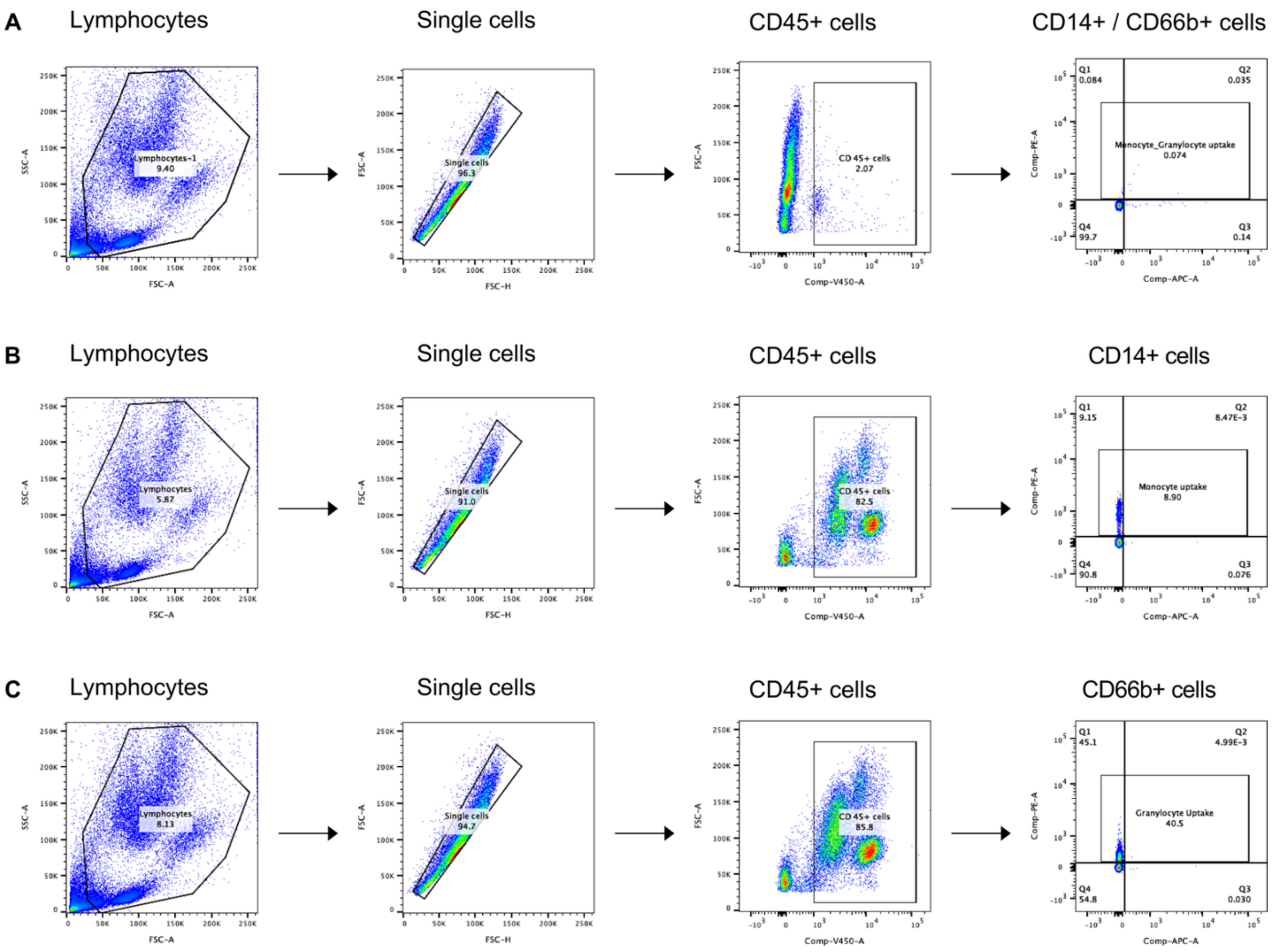


**Figure S4. Representative gating strategy for identifying monocytes and granulocytes in human peripheral blood.** Debris was excluded and leukocytes identified based on forward and side scatter (FSC/SSC) properties. Single cells were selected using FSC-A vs. FSC-H gating. $CD45^{+}$ leukocytes were then gated, followed by separation into $CD14^{+}$ monocytes and $CD66b^{+}$ neutrophils to evaluate the uptake of DiD-labeled liposomes by phagocytic myeloid cells in human blood.

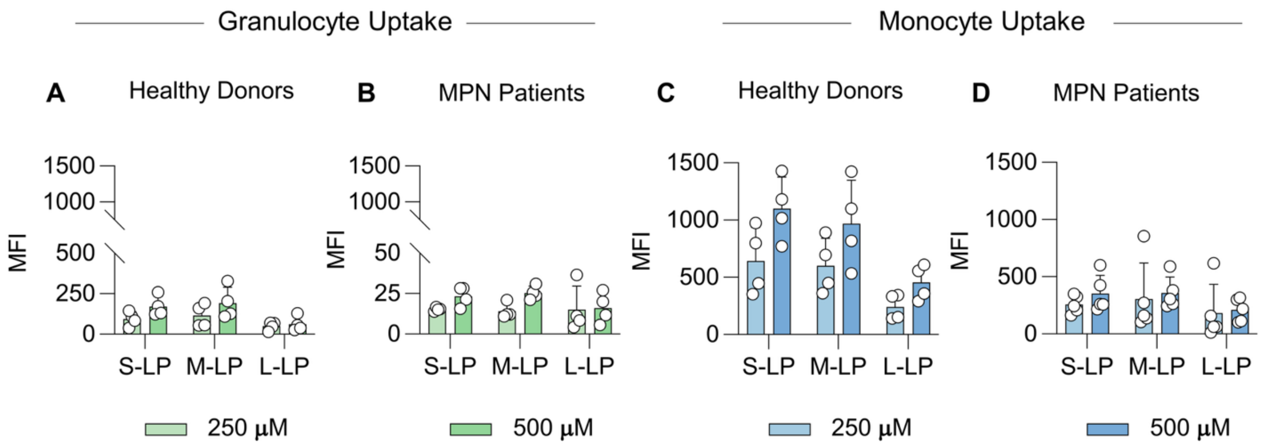


**Figure S5. LP uptake by blood myeloid cells increases with LP concentration in both HD and MPN patients.** Whole blood was incubated with LP at two concentrations (250 μM and 500 μM) for 1 h. LP uptake was quantified by flow cytometry and presented as mean fluorescence intensity (MFI) after background subtraction using unstained controls. Granulocyte uptake is shown for (**A**) HD and (**B**) MPN patients, while monocyte uptake is shown for (**C**) HD and (**D**) MPN patients. Data are presented as mean ± SD (n = 4-5).

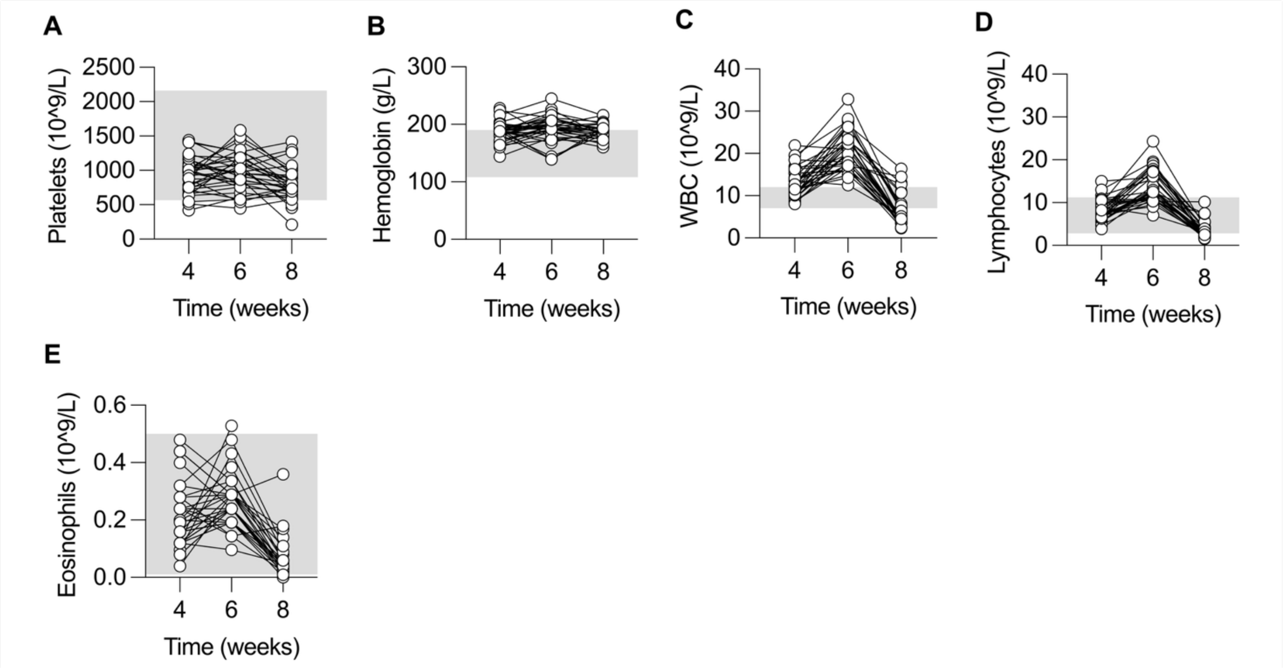


**Figure 6. PB analysis confirms PV-like phenotype in a *JAK2*$^{V617F}$ MPN mouse model.** PB was collected, and complete blood counts including (**A**) platelets, (**B**) hemoglobin, (**C**) white blood cells (WBC), (**D**) lymphocytes, and (**E**) eosinophils were measured at 4, 6, and 8 weeks post transplantation. Values exceeded the normal range reported for healthy C57BL/6 mice (grey area), indicating successful disease development.

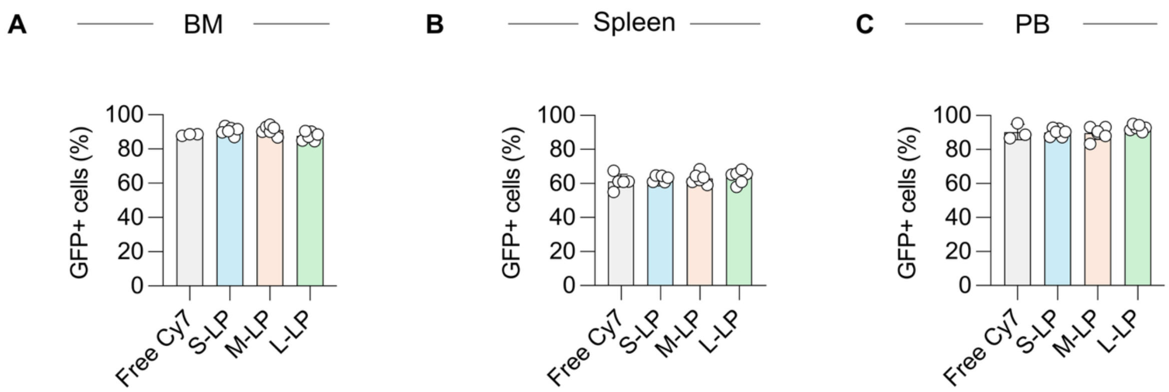


**Figure S7. Distribution of malignant GFP⁺ clones in hematopoietic organs and peripheral blood.** Flow cytometric analysis of GFP⁺ malignant cells in (**A**) BM, (**B**) spleen, and (**C**) PB showing comparable disease burden among experimental groups.

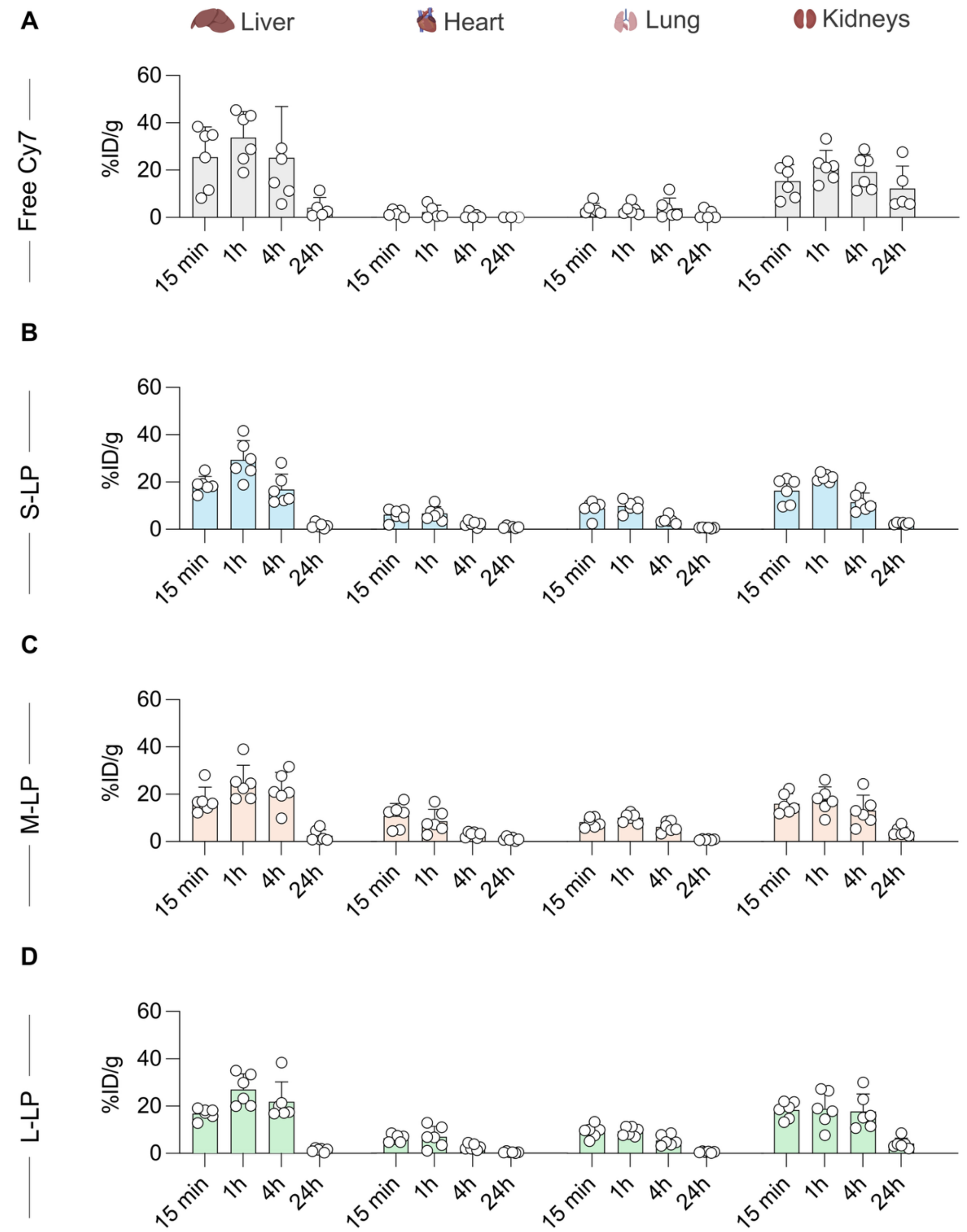


**Figure S88. Quantitative biodistribution of Cy7-labeled LP in non-target organs of the *JAK2*$^{V617F}$ MPN murine model.** LP distribution (%ID/g) following i.v. administration of (**A**) free Cy7, (**B**) S-LP, (**C**) M-LP, and (**D**) L-LP in non-target organs including the heart, lungs, liver, and kidneys. Results are given as mean ± SD; with n = 6 per group.

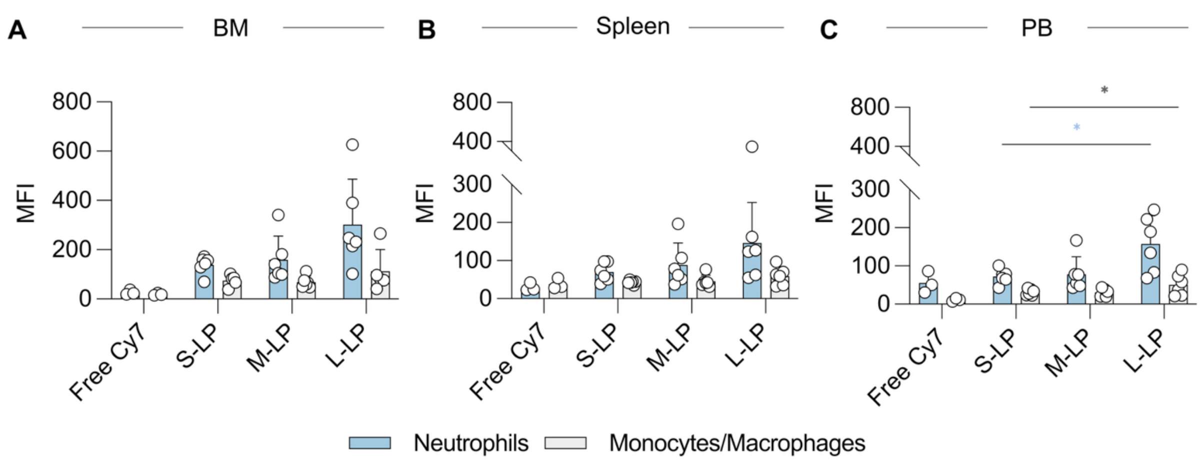


**Figure S9. Liposome size influences uptake by neutrophils and monocytes/macrophages across hematopoietic tissues.** Cellular uptake of the different LP formulations (S-LP, M-LP, and L-LP) by neutrophils (blue) and monocytes/macrophages (grey) in the (**A**) BM, (**B**) spleen, and (**C**) PB of *JAK2*$^{V617F}$ MPN mice as measured by flow cytometry 24 h after i.v. administration. LP uptake is quantified as mean fluorescence intensity (MFI) of Cy7, and results are expressed as mean ± SD (n=5-6). P values are indicated as * < 0.05, ** < 0.01, *** < 0.001, and **** < 0.0001.

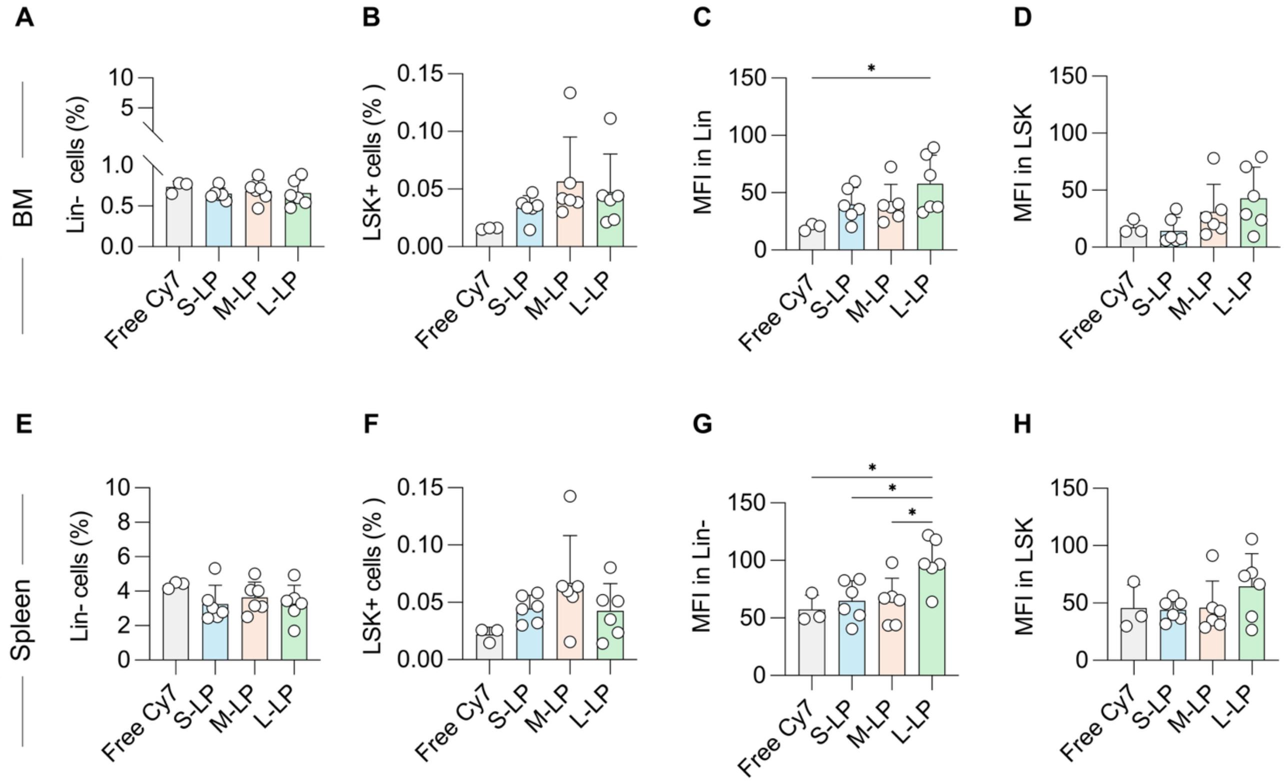


**Figure S10. Quantification of LP accumulation in Lin⁻ and LSK⁺ cell populations from bone marrow and spleen.** The frequency of (**A,E**) Lin⁻ cells and (**B,F**) LSK cells in BM and spleen, respectively. LP uptake, expressed as mean fluorescence intensity (MFI), in (**C,G**) Lin⁻ cells and (**D,H**) LSK⁺ cells from BM and spleen, respectively. Minimal LP uptake was observed in both progenitor populations across formulations. Results are expressed as mean ± SD; with n = 6 per group.
